# Bioelectronic Wearable Sensors Produced by Computerized Embroidery Using a Dual Thread Approach

**DOI:** 10.64898/2025.11.25.690465

**Authors:** Taliya Weinstein, Xintong Li, Hasan Kurt, Selin Olenik, Alexander Silva-Pinto Collins, José Manuel Rodrigueiro Flauzino, Muhammed Adeel, Laura Gonzalez-Macia, Meral Yüce, Firat Güder

## Abstract

Electronic textiles offer a promising route to wearable systems capable of continuous biophysical and biochemical monitoring. Their broader adoption remains limited by conductive threads that rarely support computerized embroidery (CEmb), reliable electronic integration, and reduced electrochemical capabilities due to coatings that degrade under bending, friction, or aqueous exposure. In this work, we use a dual-thread strategy that integrates silver-plated polyamide threads (AgPT) for high-conductivity pathways with PECOTEX, a chemically resilient PEDOT:PSS-coated cotton thread optimized for biochemical sensing. This approach preserves compatibility with CEmb, enhances electrochemical performance, and reduces cost by reserving AgPT for critical traces. We demonstrate a CEmb-compatible fabrication process for two wearable devices: a fully embroidered SmartBra for electrocardiography and a two-electrode ion sensor for potassium and sodium detection. We validate the design of our wearable dual-thread biosensors through laboratory characterization experiments and evaluate electrochemical sensing performance using a breast milk substitute, enabling realistic testing within women’s health and infant-nutrition contexts. By focusing on scalable fabrication and meaningful applications, this work advances embroidered electronic textile systems toward practical, personalized healthcare monitoring.

## 1. Introduction

Electronic textiles (e-textiles) offer a versatile platform for non-invasive wearable sensors. They integrate comfortably with the human body due to both their flexibility and conformability, making them especially valuable for health monitoring ^[1]^. Although many promising demonstrations of e-textile integration have emerged over the past decades (such as in electrocardiogram monitoring, strain, and pressure sensing), commercial adoption remains limited. Key barriers for the adoption of e-textiles include: (i) lack of compatibility with mass manufacturing techniques such as computerized embroidery (CEmb), (ii) robust chemical or biosensing applications (most notably in the durability), (iii) cost and measurement versatility ^[2,3]^. CEmb stands out among these barriers as both a scalable opportunity and a source of unique challenges. CEmb uses computer-controlled machines to stitch high-resolution (∼0.25 cm²) patterns at high speed (∼1200 stitches/min) onto various woven and non-woven fabrics, such as cotton and polyester ^[4,5]^. E-textiles benefit from the throughput and precision of CEmb, enabling highly specific designs to be reproduced seamlessly ^[6]^. Despite these advantages of versatility and high throughput, CEmb-compatible conductive threads remain a bottleneck.

The challenge of CEmb-compatible conductive threads stems from the mechanical harshness in CEmb processes. Threads are repeatedly bent and passed through a 0.8 mm needle while under tensile and frictional forces of 1–1.5 N ^[7,8]^. While conventional threads withstand embroidery without visible damage, conductive-coated threads often lose conductivity due to shearing of their outer layers ^[9]^. Even when conductivity is preserved post-embroidery, as with plated threads, electrical sensing can have mechanical challenges, such as changing base resistance over multiple stretch cycles^[10]^. Additionally, electrochemical sensing remains challenging, since most applications require electrodes with electrochemically stable coatings that are insufficiently robust to endure embroidery stresses, limiting long-term reliability ^[11]^.

For large-scale CEmb production of electrical and electrochemical sensors, the ideal conductive thread should combine structural robustness, electrochemical stability, and cost-effectiveness. At the same time, for electrochemical applications, the thread should be chemically resilient, particularly to oxidation, and exhibit either a high surface area or high electrical conductivity to ensure efficient electrochemical transduction. Finally, for practical deployment and widespread adoption, the thread must also be affordable and readily accessible. Often, no single thread satisfies all three needs. Instead, a hybrid approach may prove more practical. For example, copper threads offer excellent conductivity and moderate CEmb durability ^[12]^. Copper threads, however, are not suitable for electrochemical sensing as they are easily oxidized in aqueous conditions, and the protective coating on copper threads limits the area of the sensing surface ^[12,13]^. Silver-based (Ag) threads, such as the Ag-coated Shieldex HC-40, offer low resistance (∼114 Ω/m) and are commercially available (although not at low cost) ^[14]^. Until recently, however, Ag-coated threads could only be embroidered with highly specialized CEmb systems and were incompatible with most domestic and industrial machines ^[15]^. Ag can also be an important material for applications in wearable chemical sensing, acting as low-resistance conductors and pseudo-reference electrodes for electrochemical sensing. The electrical and chemical properties of Ag-based threads, however, degrade rapidly when subjected to electrochemical reactions ^[16]^.

To overcome these electrochemical limitations, Ag-plated polyamide threads (AgPT), such as HC-40, can be combined with the recently reported low cost PEDOT:PSS-coated cotton threads (PECOTEX), which are chemically durable and effective for sensing ^[17]^. This dual-thread approach pairs silver’s conductivity with PECOTEX’s stable sensing interface, delivering CEmb compatibility alongside reliable sensing performance. Additionally, the hybrid design enhances cost-efficiency by reserving the more expensive Ag threads for essential high-conductivity connections. Beyond retaining electrical properties through embroidery, a persistent challenge for e-textiles lies in achieving robust electrochemical sensing. Most reported platforms rely on polymer or enzyme coatings to confer selectivity for analytes such as sodium, ammonium, lactate, and pH, but these coatings frequently degrade under bending, friction, or aqueous exposure, resulting in signal drift and reduced stability during continuous wear ^[18,19]^.

In this work, we present a CEmb-compatible dual thread strategy for fabricating embroidered electrochemical and biophysical sensors using AgPT and cotton-based PECOTEX conductive threads. By pairing materials with complementary properties and refining electrode design, we address key challenges of CEmb durability and sensing stability. To demonstrate the versatility of this platform, we integrate embroidered sensors into garments for (i) electrocardiogram (ECG)-based cardiac monitoring and (ii) ionic detection of potassium (K⁺) and sodium (Na⁺) in a breast milk analog, metrics directly relevant to maternal and infant health. We further introduce a solder- and adhesive-free method for connecting sensors with flexible, battery-powered electronics, enabling seamless, continuous physiological monitoring. Importantly, our approach goes beyond prior work by directly fabricating embroidery-compatible conductive threads tailored for electrochemical applications, allowing stable and selective measurement of physiologically relevant ion concentrations.

## 2. Materials and Methods

### 2.1. PECOTEX Fabrication

We fabricated PECOTEX using a previously optimized roll-to-roll fabrication methodology ^[17]^. The thread coating formulations and the roll-to-roll fabrication setup are presented in **Figures S1 and S2**, respectively. Prior to roll-to-roll fabrication, we pretreated the virgin cotton threads (JBS Olivia, South Korea) with a desizing solution of 5 wt% Synthrapol detergent solution (Metapex 38, Kemtex, UK) for 2 h and later treated the threads with a 4:1 aqueous bleach solution for 5 h to remove waxy organic content. Finally, we washed the desized threads several times with deionized (DI) water and dried them at 50°C for 1 h. Later, we submerged the desized threads consecutively into four identical dyeing solutions with a speed of 16.5 cm·min−1 (9.9 m·h−1) and dried the threads under hot air stream before rolling them into the final bobbin. Each dyeing solution consisted of 50 mL aqueous PEDOT:PSS dispersion (1.0 – 1.3 wt% solid content, Clevios™ PH1000, Heraeus, Germany) with the additives of 3 wt% ethylene glycol (EG, Honeywell, USA) and a crosslinking agent of 5 wt% divinyl sulfone (DVS, Sigma Aldrich, Germany). We confirmed the covalent bond formation between cotton-DVS-PEDOT:PSS using Fourier transform infrared spectroscopy (Nicolet iS50, Thermo Scientific, USA) in **Figure S3**.

### 2.2. Embroidered Sensor Production

We used embroidery pattern design software (The Premier+™ 2) to create sensor electrode patterns. We transferred these designs for production to an embroidery machine (Designer Diamond Royal, Singer, Husqvarna Viking, Sweden). The base material selection was set to 95% cotton or above to match the cotton core of PECOTEX. We selected a topstitch sewing machine needle of size 80/1c2 (Schmetz GmbH, Germany) since its larger eye size prevented excessive shearing of the PECOTEX threads. Additionally, we employed a 1.8-oz tear-away stabilizer to support the stitch and prevent pattern distortion. To determine the practical feasibility of scaling up the production of the embroidery patterns, we also tested the PECOTEX thread on an industrial embroidery machine (PR1050X, Brother, USA). We produced the selected patterns on the industrial-scale machine and the standalone embroidery machine using PECOTEX and the commercially available conductive threads (Ag-plated polyamide, HC-40, Madeira Garnfabrik, Germany). Later, we measured the DC resistances of these selected patterns in both thread types using a source-measure unit (2450, Keithley, USA). A depiction of the thread bobbins and alternative sensor designs is presented in **Figures S4** and **S5**, respectively.

### 2.3. Embroidered Sensor Design and Measurement

For ECG sensing electrodes, we fixed the pattern size to 20 × 8 mm^2^, which we found to be sufficient for ECG extraction ^[20]^. For the three-terminal electrochemical electrodes, we designed the patterns to be of a similar size (25 mm × 7 mm) to that of a disposable screen-printed electrode. The working electrode had a comparable area to the screen-printed electrode, with a size ratio of 1:1.5 to the counter. The larger counter electrode was in line with other embroidered electrochemical sensors ^[21,22]^. We electrically isolated the electrode and interface contacts with an insulating silicone coating (10 mm long) to prevent wicking of the electrolyte solution. We measured the DC resistances of the embroidered patterns by conducting a four-point resistance measurement technique using a source-measure unit (2450, Keithley, USA). We performed scanning electron microscopy (SEM) using a Leo Supra 35VP instrument (Carl Zeiss, Oberkochen, Germany) to acquire micrographs of cotton and PECOTEX threads. We mounted samples directly onto conductive carbon adhesive tapes without additional preparation. Imaging was carried out under an electron beam with an accelerating voltage of 6 kV and a working distance of 6 mm. First, we embroidered sensors, and subsequently unstitched, with four-point resistance measurements obtained before and after to assess conductivity loss due to the embroidery process. We then evaluated the robustness of the design against washing abrasion over three cycles, in which we placed each embroidered sensor in a meshed washing bag and subjected it to laundering, followed by two-point resistance measurements after each cycle. Finally, we examined resistance changes of three samples of each embroidery pattern under varying embroidery parameters using an industrial machine. At a constant stitching speed of 400 stitches/min, we varied the tension of the thread from low to high (parameters summarized in **Tables S1 and S2**) and then measured the two-point resistance of the electrode designs. We then adjusted the speed and tension in combination to encompass both low (400 stitches/min) and high (1000 stitches/min) speeds under low and high tension when embroidering a basic zigzag pattern, with two-point resistance measurements collected after each condition.

### 2.4. Electrocardiogram Circuit Design and Quantification

We designed the electronic circuit to incorporate an ESP-32-WROOM microcontroller (with integrated antenna, USB compatibility, open-source voltage regulation) and an AD8232 single-lead ECG front-end chip in a standard cardiac monitoring configuration. Signal conditioning consisted of a second-order 0.5 Hz high-pass filter and a 40 Hz second-order Sallen-Key low-pass filter, yielding a 0.5–40 Hz bandwidth appropriate for single-lead ECG acquisition. The flexible printed circuit board (**Figures S6 and S7**) measures 44 × 39.2 mm and uses 18 μm copper traces, with a minimum trace spacing of 0.6 mm and teardrop-shaped terminations at all copper junctions to reduce mechanical stress. We encapsulated the flexible printed circuit board in a 6 mm layer of Dragon Skin 30 silicone rubber by creating a rectangular mold and pouring the silicone over the circuit. We then allowed the silicone to cure overnight. We attached both the circuit and a 500 mAh, 3.7 V lithium-ion battery using snap clips to the garment with the standard plier press for snap clip attachment. Coding of the microcontroller was performed in the Arduino IDE, with postprocessing and storage performed in MATLAB. We applied a moving average filter to reduce computing requirements and to process ECG signals.

We evaluated the functionality of the ECG of the SmartBra on one lab volunteer by recording 30-second intervals under four movement conditions: no movement (including breath-holding), normal breathing, swaying, and walking. We repeated each condition across three trials. Informed consent of the participant was obtained prior to data collection. We used the resulting average of signals to extract common features used to assess signal quality, such as kurtosis and signal-to-noise ratio (SNR).

### 2.5. Electrochemical Ion Detection

The two-terminal working electrode comprised two parts: a 7 mm PECOTEX-embroidered sensor and an 18 mm commercial AgPT connector. The AgPT connector is electrically isolated from the solution via a Dragon Skin 30 silicone coating, applied with a spatula on both sides of the material and allowed to cure overnight. We coated the PECOTEX sensor with either a K⁺ or Na⁺ selective membrane cocktail, following the composition designed by Gao *et al.* ^[23]^. All chemicals are purchased from Sigma-Aldrich/Merck unless otherwise stated, with code numbers provided after the mentioned reagents. respectively

We fabricated a Na^+^ ion-selective membrane using the following Na^+^ ionophore cocktail consisting of 1 wt% sodium ionophore X, 0.55 wt% sodium tetrakis[3,5-bis(trifluoromethyl)phenyl] borate, 33 wt% high molecular weight polyvinyl chloride, and 65.45 wt% bis(2-ethylhexyl) sebacate. We dissolved 50 mg of Na^+^ ionophore cocktail in 300 µL of tetrahydrofuran solvent to modify the embroidered electrodes. Similarly, we fabricated a K^+^ ion-selective membrane using the following K^+^ ionophore cocktail consisting of 2 wt% potassium ionophore I, 0.5 wt% sodium tetraphenylborate, 33 wt% high molecular weight polyvinyl chloride, and 64.5 wt% bis(2-ethylhexyl) sebacate. We later dissolved 100 mg of K^+^ ionophore cocktail in 350 µL of cyclohexanone solvent to modify the embroidered electrodes. Electrodes were immersed in the ionophore solution for 10 minutes and dried in a fume hood for 3–4 h, and the procedure was repeated once to ensure uniform coverage. We used the commercial AgPT for embroidering a 7 mm reference electrode, electrically isolated from the connector in the same manner as for the working electrode (described above). We used a potentiostat (EMStat 3, PalmSens, Netherlands) for cyclic voltammetry measurements on the three-electrode sensor configurations and for open-circuit potential (OCP) measurements in the two-electrode setup, where OCP reflects the electrode’s intrinsic electrochemical stability under no applied current.

We prepared a Tris buffer solution by titrating with HCl. For calibration in Tris buffer, we immersed sensors in analyte solutions for 1 minute before recording. Each OCP trace lasted 100 seconds, with potentials acquired every 0.2 seconds in a 10 mL volume. Measurements were taken from high to low concentrations (e.g., K⁺ from 100 mM to 0 mM), starting with 100 mM KCl (P9541) or 30 mM NaCl in Tris (S9888) for the K⁺ and Na⁺ ionophore-coated working electrode, respectively, and reducing concentration by serial dilution.

We used the same measurement approach in Aptamil, diluted with deionized water to yield baseline concentrations of 0.3 mM Na⁺ or 1 mM K⁺. The initial solutions contained either 100 mM KCl or 30 mM NaCl in diluted Aptamil, with subsequent concentrations obtained by stepwise dilution to 1 mM K⁺ or 0.3 mM Na⁺.

To assess selectivity, we first measured responses in Tris (0 mM analyte) and then in 5 mM analyte solutions prepared by adding 500 µL of 100 mM analyte stock to 9.5 mL Tris. The following analyte solutions were tested: KCl (P9541), KH_2_PO_4_ (P5379), KNO_3_ (P8291), NH_4_Cl (21330), MgCl (M8266), NaCl (S9888), NaH_2_PO_4_ (567547), and NaNO_3_ (221341). Sensors were immersed for 1 minute before measurements started. OCPs were recorded for 100 seconds at 0.2-second intervals, and electrodes were rinsed with deionized water between analytes. Measurements were baseline-corrected by subtracting the recorded 0 mM potential.

We evaluated stability by recording OCPs every second during continuous immersion. For the 24 h test, three working electrodes were connected to an MUX8-R2 Multiplexer (PalmSens, The Netherlands) and measured simultaneously against a single reference (AgPT) for 86,400 seconds.

Following stability experiments, we recalibrated the sensors in Tris and Aptamil using the same procedure described above. Three electrodes were connected through a MUX8-R2 Multiplexer (PalmSens, The Netherlands) with one reference electrode (AgPT) and immersed in 10 mL of analyte solution for 1 minute before OCP measurement.

For the three-electrode design, the working electrode comprised a 7 mm PECOTEX-embroidered sensor, connected to an 18 mm commercial AgPT connector, which is electrically isolated from the solution via a Dragon Skin 30 silicone coating for improved conduction. The reference electrode had the same design yet was manufactured completely from the AgPT, while the counter had a length of 1.5x the size of the working electrode’s length. For the scan rate study, we submerged the three-electrode sensor in 0.1 M KCl as we increased the scan rate from 20 mV/s to 350 mV/s. For the stability study, we fixed the scan rate at 100 mV/s while we submerged the sensors in 0.1M KCl over 90 scans. For both, we used a potential window from -1.2 V to 1.2 V, with the recordings made on a potentiostat (EMStat 3, PalmSens, Netherlands).

## 3. Results and Discussion

### 3.1 Design Considerations of the SmartBra

The final version of our integrated e-textile platform, the “SmartBra,” incorporates thread-based ECG electrodes and ion-selective electrochemical sensors into an off-the-shelf sports bra using CEmb (**Figure 1A**). We select these two sensing modalities to demonstrate how embroidered e-textiles can capture both biophysical and biochemical signals relevant to postpartum monitoring. Maternal and infant health remain areas with notable measurement gaps, in part due to the historical underrepresentation of women in medical research^[24,25]^. Expanding access to comfortable, on-body sensing interfaces may help address these gaps by enabling the routine collection of clinically meaningful signals. Breast milk is a clear example: it plays a critical nutritional role in the first six months of life, with deficiencies in nutrients and bioactive compounds impairing infant growth ^[26]^. Its composition, however, is typically assessed only through laboratory testing, limiting timely feedback for mothers. Continuous, on-body monitoring could offer more immediate insights and support more adaptive infant nutrition ^[27]^.

**Figure 1.**
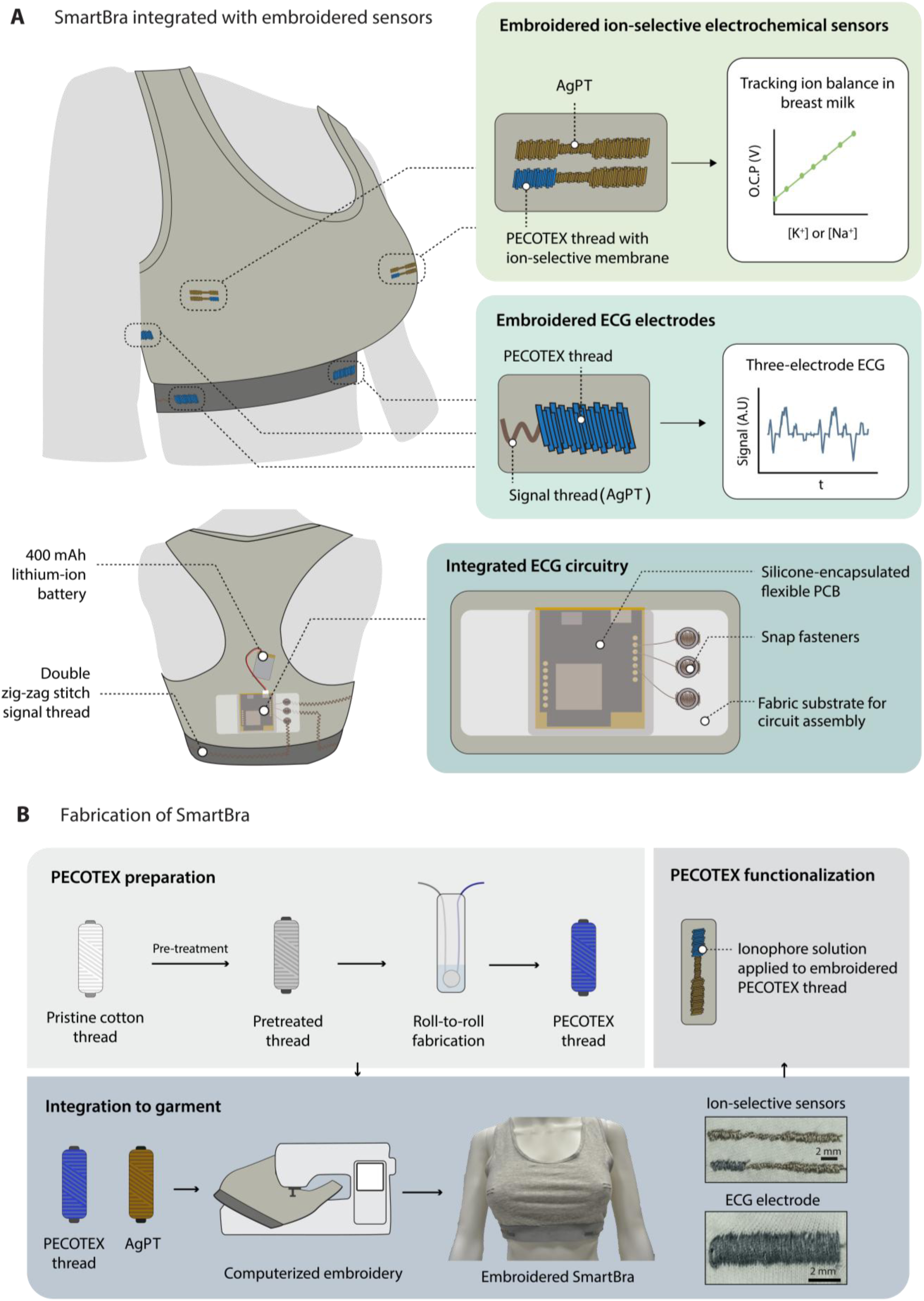
SmartBra design overview and key fabrication steps. (**A**) A schematic illustration of the SmartBra design, with ECG sensing located in the elastic bands and the ion-based breast milk sensing over the nipple area for passive milk collection. (**B**) Schematic illustration of the fabrication of the SmartBra using embroidery of PECTOTEX and AgPT (Ag-plated thread). Roll-to-roll PECOTEX production (above) and subsequent computerized embroidery (bottom).

To produce this multimodal sensing platform, we utilized both cotton-based PECOTEX and commercially available AgPT, enabling biomedical/electrochemical signal monitoring and robust low-resistance interconnects. We positioned the electrochemical electrodes over the nipple-areola complex (NAC) region to track the ionic content of milk discharge in lactating women. Specifically, we aimed to track the Na^+^/K^+^ ratio in breast milk, which is highly important to both lactation assessment and the health of the infant ^[28]^. The electrodes consist of satin stitches, which provide a continuous surface area for ion tracking. OCP is used to measure the changes in ion concentration, as potentiometric measurements proved more robust for the high-resistance PECOTEX thread.

We investigated several locations on the garment to optimize ECG signal quality and found the elastic band region (below the chest) to be ideal for ECG monitoring as it secures reliable skin contact for a range of motions. Other anatomical locations tested (see **Figure S8**) exhibited greater susceptibility to motion artifacts and intermittent signal loss, consistent with previous reports ^[29]^. The ECG electrode consists of a wide satin stitch (continuous surface with high thread coverage), which is connected to the electronics with AgPT in a double zigzag stitch. We chose the zigzag pattern because it enables the material to stretch easily while the bra is deformed without affecting the stitch integrity. To fabricate the SmartBra, we needed to transfer a high-redundancy embroidery pattern from a digital design to the computerized embroidery system (**Figure 1B)**. The process starts with the PECOTEX production using the same procedure as in Al Shabouna *et al* ^[17]^. We then used the PECOTEX thread for the working electrode of the two-terminal electrochemical sensors and the ECG satin stitch electrodes. Functionalization of the PECOTEX for electrochemical sensing occurs post-embroidery to ensure the ionophore coating does not get sheared off under the mechanical forces of the embroidery process.

To enable a fully functional and integrated embroidered sensor, we optimized the embroidery process to limit degradation. We designed the flexible circuit to be comfortable with body movements, while the encapsulation layer protects the circuit from warping. Interfacing rigid electronic components with conformable e-textiles remains a substantial challenge due to mechanical mismatches and requirements for washability, necessitating solutions that allow secure yet detachable connections. We secured the circuit together with the battery to the bra with snap fasteners, which act as a clip-on conductive connection, enabling easy removal for washing.

### 3.2 Electrode and Thread Characterization

We investigated different electrode embroidery patterns for ECG and electrochemical monitoring to determine which design best preserved the functional integrity of the PECOTEX thread post-embroidery (**Figure 2A)**. The key determinants of the retention of electrical performance included stitch density and the number of times the PECOTEX thread penetrated through the material ^[30]^. These factors must be balanced against excessive PECOTEX re-embroidering, as more thread will invariably lead to better electrical conductance because more parallel lines are created ^[14]^. We evaluated two embroidery patterns: the common e-textile square-filled crosshatch ^[17]^ and a satin stitch. We chose the crosshatch design because it is approximately 70% more thread-efficient, requiring a shorter thread length. We selected the satin stitch because it preserves a continuous conductive path in response to the mechanical abrasion from embroidery. In addition, the satin stitch offers greater stitch density and thus a larger surface area, consistent with prior recommendations to maximize active sensing regions ^[31]^. We designed the electrochemical sensor to comprise a pseudo-reference electrode made from AgPT and a working electrode with the top 7 mm constructed from PECOTEX. We used AgPT for all remaining connections fabricated to enhance conductivity and ensure robust conductive connections. We embroidered both the working and reference electrodes of the electrochemical sensor with a satin stitch, as it offered a consistent surface area for stable reaction kinetics and reliable signal sensitivity. Additionally, the satin stitch demonstrated superior reproducibility and mechanical durability (evaluated in terms of conductivity changes) compared to alternative patterns (alternative designs in **Figure S9**).

**Figure 2.**
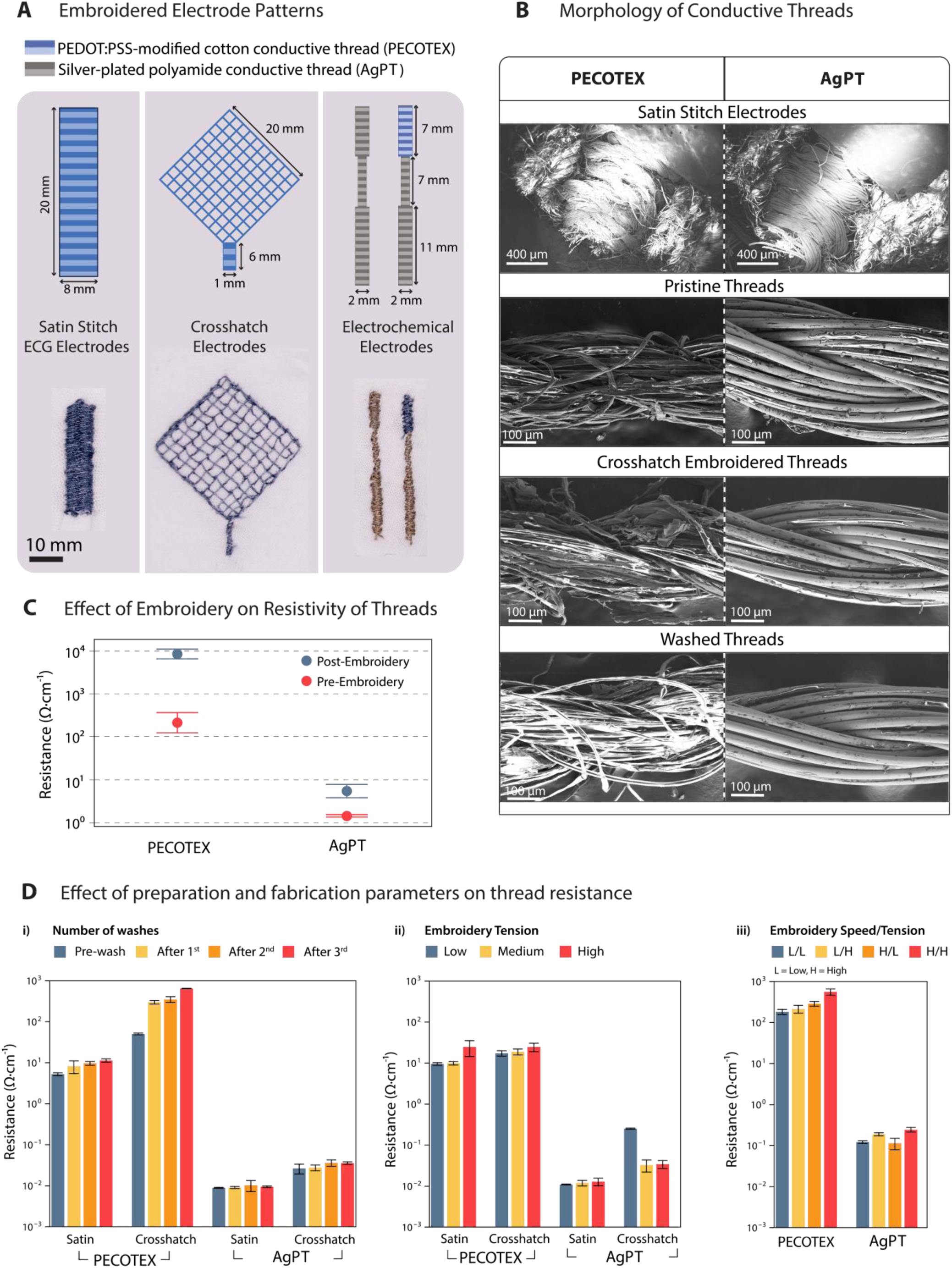
Characterization of the embroidered electrode design and thread under different mechanical stresses (**A**) Design and realized configurations of ECG and electrochemical electrodes. Crosshatch and satin stitches were tested for ECG electrodes, while electrochemical electrodes used only satin stitch. The shades of grey represent commercial AgPT, while the shades of blue represent the PECOTEX threads. (**B**) SEM images of the satin-stitched electrode (top) and of both PECOTEX and commercial AgPT under different conditions, with the same scale. (**C**) The resistance per meter length of thread for commercial AgPT and PECOTEX thread pre- and post-embroidery (n = 5). A basic crosshatch pattern was embroidered and then unstitched for the after-embroidery measurement. (**D**) The effect of (**i**) machine washing, (**ii**) thread tension, and (**iii**) embroidery speed on the resistances of both PECOTEX and AgPT with satin- and crosshatch stitches after embroidery on an industrial machine.

To evaluate the microstructural changes to the sensors and threads, we performed scanning electron microscopy (SEM) after mechanical abrasive procedures, such as embroidery and washing (**Figure 2B**). Post-embroidery, the thread from the satin stitch pattern experienced minimal fraying and maintained a consistent conductive area in both the PECOTEX and the AgPT. The resilience of the satin stitch can be attributed to the redundant conductive paths produced by the multiple overlapping threads in this pattern, enabling sensor functionality even when individual threads are damaged. Conversely, we observed evident thread degradation in the crosshatch pattern, likely due to the increased penetration through the material. These abrasive forces of both penetrating the material and the embroidery process partially stripped the conductive coating from PECOTEX threads and caused delamination in commercial AgPT (see **Figures S10, S11,** and **S12**). Post-wash SEM inspection further confirmed material degradation, with PECOTEX exhibiting more severe fraying than the AgPT, likely due to the stronger bonding resulting from the silver-plating manufacturing process on the polyamide thread.

To quantify the conductivity degradation of both threads, we performed resistance measurements before and after embroidering a standardized 20 mm² crosshatch pattern (**Figure 2C**). We then unstitched the pattern and compared its resistance to pristine unembroidered thread. The pristine unembroidered PECOTEX had a resistance of 245 ± 121.76 Ω, two orders of magnitude higher than the unembroidered AgPT (1.44 ± 0.09 Ω). We expected the resistance to differ between materials due to differences in base conductivity and fabrication. Following embroidery, the AgPT showed a modest 4-fold resistance increase to 5.78 ± 1.99 Ω, suggesting that the plating offers protection against needle-induced abrasion. In contrast, PECOTEX exhibited a 35-fold increase in resistance (8736.88 ± 2236.11 Ω), attributed to dye coating loss during stitching (see **Figure S10** for further detail). These results highlight the importance of selecting appropriate thread materials and stitch patterns to optimize sensor durability and performance.

To assess the real-world robustness of PECOTEX sensors, we evaluated their electrical performance following machine washing (**Figure 2Di**). We tested three samples of each embroidery pattern, 8 mm satin stitch and 2 mm cross-hatch, for baseline and post-wash DC resistance. The satin stitching exhibited the lowest initial resistance (1006.66 ± 73.71 Ω) due to only the border of the pattern being damaged by material piercing. In contrast, the crosshatch pattern displayed a higher resistance (4-fold increase) and greater variability, likely due to more frequent thread piercing and associated dye shearing. After three wash cycles, only one of the crosshatch electrodes remained measurable, indicating critical failure in the single-threaded structure where any break disrupted the circuit. The satin stitch electrodes, with overlapping threads, retained conductivity across all samples, despite a resistance increase of 59% after the first wash and 118% after the third. The commercial AgPT patterns showed minimal resistance increases (<55%), with this thread optimized to withstand wash damage. Measurement limitations, including probe placement damaging the thread and resistance point inconsistency, should be noted.

We also investigated the industrial scalability of both AgPT and PECOTEX embroidery using an industrial embroidery machine under varying machine tensions and speeds (**Figures 2Dii and 2Diii**; parameters in **Tables S1 and S2**). At a constant speed of 400 stitches/minute, increased tension led to a rise in resistance of 1.37x in the PECOTEX satin stitch and 1.17x in the AgPT’s satin stitch (**Figure 2Dii**). We observed a 2.57x increase in the resistance of the PECOTEX crosshatch pattern, emphasizing its sensitivity to mechanical stress due to limited stitching redundancy. Despite greater degradation at the higher levels of tension for the industrial machine, PECOTEX maintained sufficient conductivity.

With thread tensions affecting the aesthetic and the ease of embroidery, tension for sensor production can be manipulated to ensure the embroidered sensors remain functional when produced at scale. The stitching speed is also critical for scalability, affecting productivity and profitability of the production site. We embroidered a wave-like pattern at low (400 stitches/min) and high (1000 stitches/min) speeds under low and high tension (**Figure 2Diii**). Increasing embroidery speed from 400 to 1000 stitches/min at high tension led to a 157.24% resistance increase in PECOTEX and 31.65% in AgPT. PECOTEX was more vulnerable overall, with two thread breakages at high speed. These results suggest that stitching speed has a more pronounced impact on conductivity than tension and must be carefully managed in industrial settings. Nevertheless, both thread types remained functional under optimized conditions, supporting the feasibility of scalable embroidered sensor production.

### 3.3 Circuit Design, Fabrication, and Embroidery Interfacing

The design of the SmartBra ensures stable acquisition of biological signals by optimizing integration of electronics, both electrically and mechanically, for motion-resilient performance and user comfort (**Figure 3A**). We designed robust and reliable thread-to-circuit connections that decrease resistance at interface (resistance decreasing by at least 38% as shown in **Figure S13**) while securing the circuit in place using a double running stitch, which begins 10 mm from the connection point, moves toward the circuit, and loops back to the starting position (**Figure S14**). We attached both the circuit and lithium-ion battery (**Figure 3A**) using snap clips, a standard solution for detachable textile electronics, to enable easy removal before washing and reduce mechanical strain during use. We positioned the circuit along the right side of the spine to minimize motion artifacts, where spinal bending exerts less mechanical stress. We stitched connections between the electrodes and the circuit using a stretchable zig-zag pattern, enabling material stretching without stitch damage ^[32]^. An equivalent AgPT bra layout is shown in **Figure S15**.

**Figure 3.**
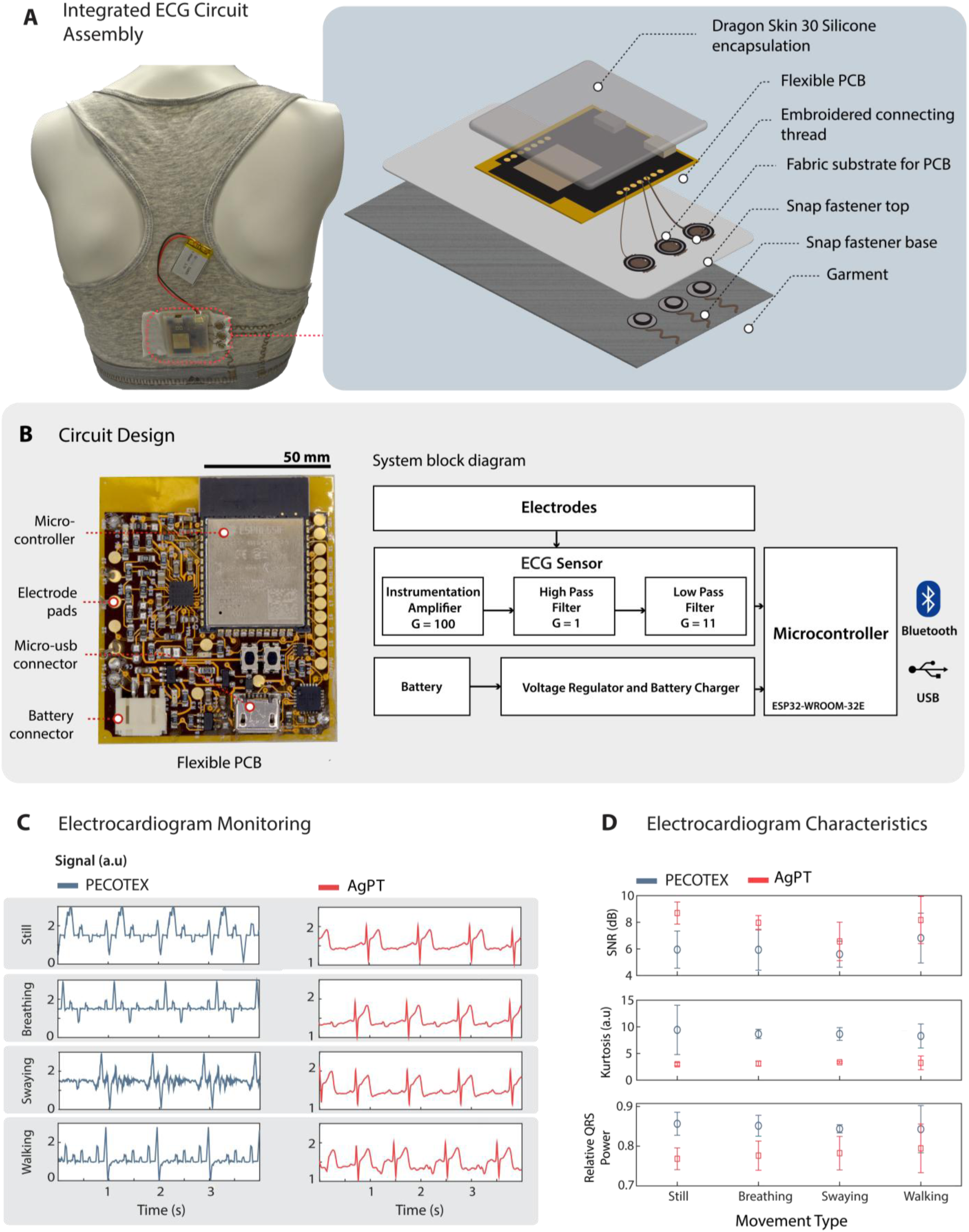
Design and characterization of the ECG monitoring circuit (**A**) Photograph of the back of the SmartBra (left), where the silicone-encapsulated circuitry (right) and sew-on snaps interconnect to the circuitry. (**B**) Photograph of flexible circuit board (left) and block functional diagram of the circuit (right), with G being the amplification gain of each of the circuit stages (**C**) ECG signal readout in various conditions (while standing still, breathing, swaying, and walking) using both PECOTEX (left blue panel) and AgPT (Ag-plated thread, right red panel). (**D**) ECG characteristics are represented in terms of signal-to-noise ratio (SNR), kurtosis, and relative QRS power.

The electronics architecture (**Figure 3B)** incorporates an ESP-32-WROOM microcontroller, which we selected for its integrated antenna that eliminates the need for external modules, reducing device size and complexity. The built-in USB compatibility allows straightforward programming and power management, while the integrated antenna ensures reliable operation with minimal additional hardware, making the system cost-effective and accessible. We chose to use Bluetooth for data transmission due to its effectiveness over long distances and lower power consumption than other wireless data transmission protocols. For cardiac monitoring, we used the AD8232 single-lead ECG front-end chip, as it offers a compact, low-power solution optimized for wearable devices. We implemented the standard single-lead ECG configuration because it provides clinically relevant heart rate signals while maintaining simplicity of integration. This configuration includes an instrumentation amplifier to extract weak, millivolt-level ECG signals ^[33]^. The amplifier’s high input impedance prevents electrode loading, while its high common-mode rejection suppresses motion and mains noise. We incorporated two analog filters to improve signal quality: a 0.5 Hz second-order high-pass filter to remove baseline drift caused by respiration and body motion, and a 40 Hz second-order Sallen-Key low-pass filter (**Figure S13**) to ensure that the signal is not saturated by high-frequency noise (i.e., mains interference). Together, this 0.5–40 Hz bandwidth aligns with established single-lead ECG standards, ensuring that key cardiac features (P, QRS, and T waves) are captured with sufficient fidelity while minimizing power consumption and avoiding unnecessary signal artifacts ^[34]^. Additionally, we employed an amplification of 1100 to bring the ECG signal within the input range of the analog-to-digital converter (ADC) of the ESP-32-WROOM microcontroller ^[33]^. Once assembled, we encapsulated the circuit in 6 mm of silicone (Dragon Skin 30 silicone rubber), a crucial step to protect both the thread-circuit interface and the circuit board itself against mechanical impact or torsion from body movements.

### 3.4. Electrocardiography

We validated the performance of the non-invasive ECG sensor through in situ monitoring continuously, recording over four movement conditions (conditions listed here, **Figure 3C**). Since movement introduces noise and motion artifacts into the signal, we wanted to demonstrate the operation of the device over no motion (still and not breathing), simple breathing, and then two different forms of displacement: swaying horizontally and walking straight ahead. The commercial AgPT bra consistently produced a defined QRS complex (0.0088 s at rest to 0.12 s while walking) and QT interval (0.33 s at rest to 0.35 s while swaying), all within normal ranges ^[35]^. In contrast, the PECOTEX bra did not record the full QRS complexes, often capturing only the R wave. Where a QRS complex was identifiable (0.152 s at rest, 0.132 s while swaying), values exceeded the upper normal physiological defined limit (0.06–0.11s) by 38% and 20%, respectively, indicating distortion and noise interference ^[35,36]^. Despite these waveform limitations, the PECOTEX bra consistently captured the R wave, enabling reliable heart rate monitoring.

We computed the signal quality indices (SQIs) from 35-second ECG recordings, including signal-to-noise ratio (SNR), kurtosis (which indicates the presence of spike-like deviations), and relative QRS power ^[30,37]^ (**Figure 3D)**. The AgPT bra outperformed PECOTEX across all metrics. Notably, SNR was significantly higher in the AgPT bra during stillness (*p* = 0.0435), while PECOTEX signals exhibited kurtosis values well above the noise threshold (>5), reflecting excessive artifact contamination. Relative QRS power for PECOTEX fell outside the expected range (0.5–0.8)^[36]^, confirming degraded signal integrity. Electrical resistance at the electrode-to-circuit interface correlated with these outcomes. AgPT connections remained below 52 Ω, while PECOTEX connections reached up to 650 kΩ. Additionally, the longer PECOTEX lead introduced a 37 kΩ resistance imbalance, impairing common-mode rejection and increasing susceptibility to interference ^[38,39]^. Increasing stitch density may help reduce resistance and improve performance ^[17]^. Although current PECOTEX-based designs exhibit limited ECG fidelity, improved stitching strategies and connection optimization could enhance their viability for low cost, embroidered health monitoring systems. Alternatively, AgPT could be employed for measurements necessitating high conductivity.

### 3.5 Electrochemical Characterization

To demonstrate the potential of the dual tread approach to electrochemical textile-based sensing, we employed functionalized electrochemical sensors incorporating PECOTEX and AgPT thread into textile-based interfaces ^[17]^. PECOTEX’s ability to support reliable electrochemical readouts highlights its suitability for wearable sensing platforms. We demonstrated this quality through the integration of OCP monitoring of K⁺ and Na⁺, key breast-milk electrolytes essential for bone formation in infants, metabolism, neurodevelopment, and muscle function ^[40,41]^. All recordings and data storage were performed using a potentiostat (EMStat 3, PalmSens, Netherlands).

The US Institute of Medicine recommends an intake of 0.4 g potassium/day for infants under six months ^[42]^ corresponding to a breast milk concentration of 11.18–15.16 mM K⁺ ^[43]^. Na^+^ is additionally essential for body growth and neurodevelopment, with hyponatremia being commonly experienced in childhood ^[44,45]^. The National Institute of Health recommends a mean concentration of 6 mM Na^+^ in breast milk for infants within the first 6 months, translating to a range of 3.85 mM to 8.32 mM ^[43]^. To ensure infants do not experience any ion deficiencies, monitoring of K^+^ and Na^+^ ion levels in lactating women is essential.

We designed a two-terminal sensor with a PECOTEX working electrode and a pseudo-reference of AgPT (7 mm x 2 mm surface area), both embroidered with a satin stitch to enable reproducible continuous thread area for the ion interaction. To enable ion sensing, we coated the working electrodes with ionophore cocktails (ion-binding molecules that selectively transport target ions across a membrane). For Na⁺ sensing, we used sodium ionophore X, and for K⁺ sensing, potassium ionophore I, each prepared in a standard polymer–plasticizer matrix with ionic additives following the same recipe as Gao *et. al.* ^[46]^. To investigate the performance of the developed ion-sensitive sensors, we first conducted experiments in a controlled solution with Tris buffer (**Figure 4A)**. Breast milk contains multiple electrolytes, so sensor selectivity is essential for accurate analyte monitoring. Cl⁻ and K⁺ are highest in breast milk (∼11 mM and ∼12 mM), with Na⁺ (∼5 mM) and Mg²⁺ (∼1 mM) also present ^[47]^. We tested the K⁺ and Na⁺ sensitivity of the Na⁺ and K⁺ sensors, respectively, to assess potential cross-interference because these ions are the dominant monovalent cations in breast milk and the most likely mutual interferents due to their similar properties ^[48]^. We recorded OCP in Tris buffer containing 5 mM non-target ions, a mid-millimolar level chosen for physiological relevance to the order of magnitude of breast-milk electrolytes. Under these conditions, non-target electrolytes caused negligible interference in the responses of K⁺ and Na⁺ sensors.

**Figure 4.**
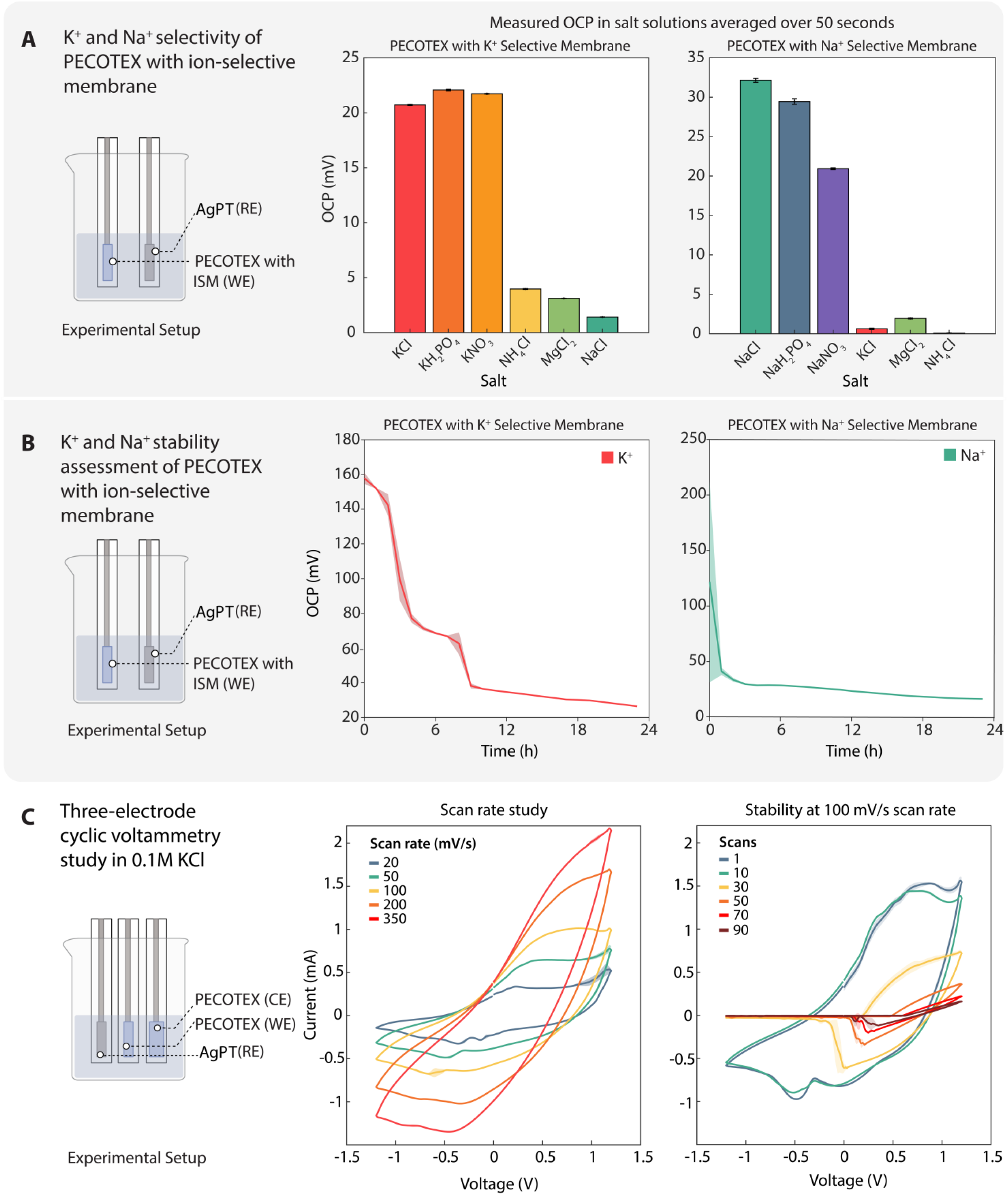
Electrochemical characterization of two- and three-electrode sensor configurations (**A**) Experimental setup for selectivity assessment of K^+^/Na^+^ ion-selective electrodes against AgPT (Ag-plated thread) reference in Tris buffer(left) and results (right); (**B**) Experimental setup for stability measurements of K^+^/Na^+^ ion-selective sensors in Tris buffer spiked with 13 mM KCl and 5 mM NaCl, respectively (left) and results (right); (**C**) Experimental setup for three-electrode sensor configuration (left) with scan rate study of the three electrode sensor from 20 mV/s to 350 mV/s in 0.1 M KCl (middle) and stability study at 100 mV/s in 0.1M KCl over 90 scans (right).

To evaluate long-term operational stability for longer-term monitoring of milk, we immersed the sensors in their respective mean breast milk ion concentrations (K⁺ sensor in 13 mM KCl and the Na⁺ sensor in 5 mM NaCl) for 24 h (**Figure 4B**). The K⁺ sensor required 10 h to reach steady state, whereas the Na⁺ sensor stabilized in 3 h. Even after 24 h in solution, a residual diffusion period occurs because ions continue to redistribute inside the thick plasticized PVC membranes, with the time to stabilization growing roughly proportional to membrane thickness squared ^[49]^. The K⁺ sensor is slower to equilibrate than the Na⁺ sensor because valinomycin (K⁺ ionophore) forms especially stable complexes with K⁺, increasing site occupancy and effectively lowering mobility in the film, whereas Na⁺-selective membranes based on Sodium Ionophore X exhibit faster transport under comparable conditions ^[50,51]^. This multi-hour equilibration is still fit-for-purpose for breast-milk monitoring because clinically relevant shifts in Na^+^ and K^+^ during lactation change over hours rather than minutes ^[52,53]^. If a faster equilibration time is required, the membrane thickness can be readily adjusted by reducing the dip duration or the number of coating cycles. Following this diffusion period, the sensor remained stable over the remaining 12 h of testing (∼1.6 mV·h^−1^ drift for K^+^, ∼0.83 mV drift for Na^+^), achieving similar stability to the literature ^[46]^

To demonstrate proof-of-concept for a three-electrode sensor configuration, we used PECOTEX thread for the working and counter electrodes and a commercial AgPT as the reference (size ratio 1:1.5:1) and for all connections (**Figure 4C)**. A three-electrode design in the SmartBra would expand sensing capabilities beyond electrolytes, supporting techniques such as cyclic voltammetry, chronoamperometry, and square-wave voltammetry for real-time biomarker tracking, including aptamer or antibody-based or enzymatic reactions ^[54]^. We evaluated the performance of the three-electrode sensor performance using cyclic voltammetry in 0.1 M KCl. We applied a potential window of −1.2 V to 1.2 V, consistent with other modified textiles ^[55]^. A scan rate study (20–350 mV/s) revealed that lower scan rates (20–100 mV/s) enabled the characteristic PEDOT:PSS double peak in PECOTEX ^[56]^.

We also evaluated the electrochemical stability of the PECOTEX working electrode using cyclic voltammetry in 0.1 M KCl (90 cycles, 100 mV/s scan rate) (**Figure 4C-right**). Although stable for the first 10 cycles, there is a marked decrease in the current amplitude after 30 cycles, which continues to rapidly decrease over the remaining 60 cycles. Since embroidery causes the polymer to shed off the sensor’s surface (**Figure S10**), we believe that the sensor reaches overoxidation sooner than similar sensors previously reported using a single thread ^[17]^. This overoxidation and polymer coating damage would damage the sensor’s structural integrity on successive cycles, limiting the electrochemical application of the embroidered sensors to a fixed number of cycles, which is sufficient for shorter-term or single-point measurements ^[57]^. While we are able to demonstrate proof-of-concept operation for shorter-term or single-point measurements, further optimization of the material and mechanical properties will be required to achieve the long-term robustness needed for wearable applications.

We conducted a three-part study to quantify sensor sensitivity and validate performance in a buffer and a breast milk-like matrix, Aptamil. First, we evaluated PECOTEX working electrodes (WE) with an Ag/AgCl reference in Tris buffer by recording OCP for 100 s at 0.2-s intervals across physiologically relevant stepped ion ranges (K⁺: 1–100 mM; Na⁺: 0.3–30 mM), shown in **Figure 5A**. To span the detection limits, concentrations were chosen at 10× above and below the physiological range. We observed the OCP traces rise with increasing concentration and remain nearly flat over 100 s after spiking, consistent with a Nernst-type behavior and good short-term stability. Second (**Figure 5B**), we tested the same protocol with a commercial AgPT as the WE in Tris to highlight the necessity of hybrid fabrication with PECOTEX for electrochemical applications. The AgPT WE, as expected, failed to yield a usable calibration due to poor adhesion of the ionophore on the hydrophobic yarn. Finally, we conducted an experiment measuring OCP in Aptamil using an embroidered PECOTEX WE and an AgPT pseudo-reference (**Figure 5C).** We noted that in Aptamil, the sensor demonstrated a flatter OCP–time trace compared with the Tris buffer solution (**Figure 5A**). This behavior arises because infant-formula matrices such as Aptamil have higher, more constant ionic strength (Aptamil = 310 mOsmol/L ^[58]^). To contextualize sensor performance, we plotted the respective calibration curves of the abovementioned experiments (**Figure 5D**).

**Figure 5.**
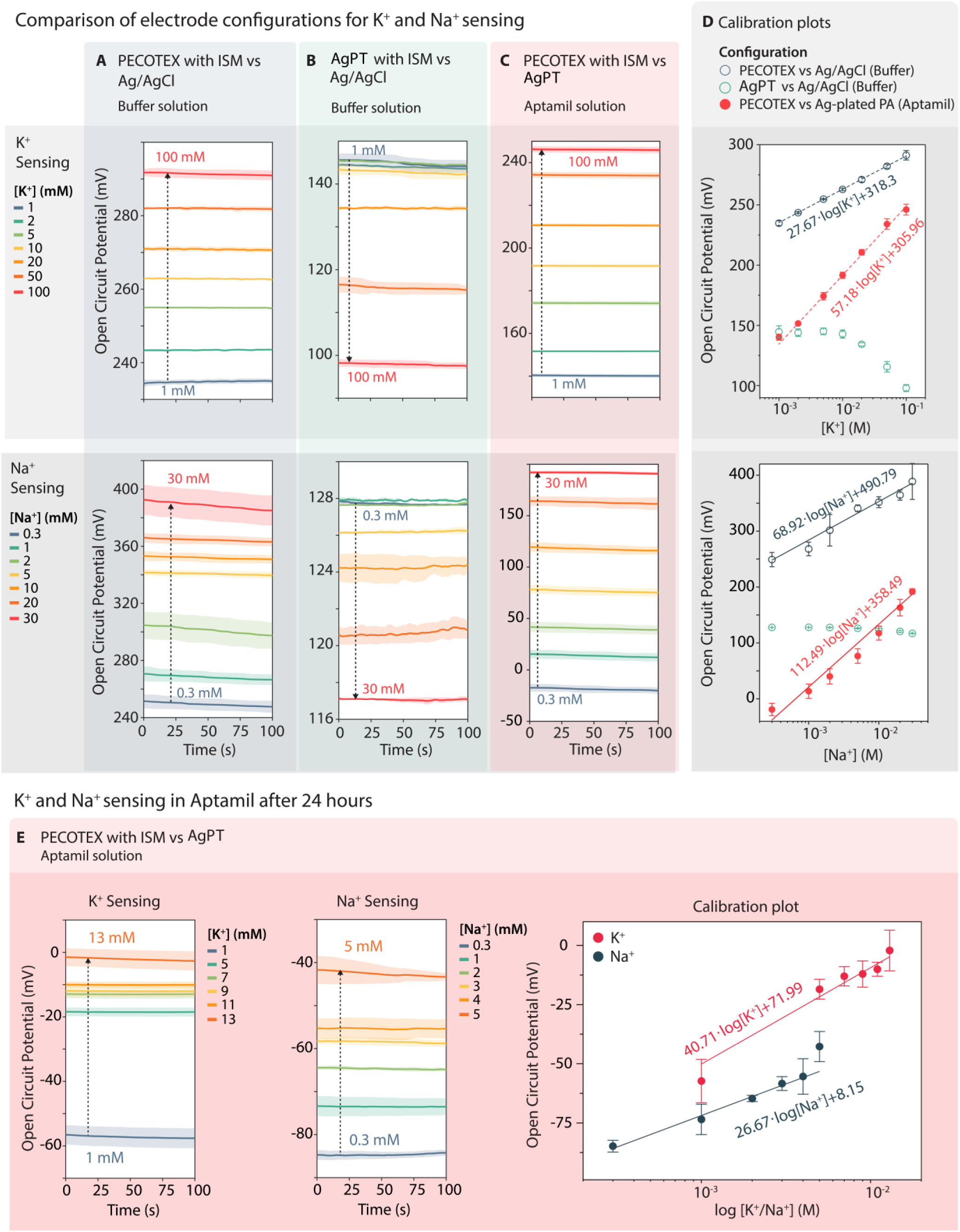
OCP calibration results for different ion-sensitive (K⁺/Na⁺) sensor configurations. **(A)** Sensor configuration using PECOTEX as the working electrode and commercial Ag/AgCl as the reference electrode, tested in Tris buffer. **(B)** Sensor configuration using AgPT (Ag-plated thread) as the working electrode and commercial Ag/AgCl as the reference electrode, tested in Tris buffer. **(C)** Sensor configuration using PECOTEX as the working electrode and AgPT as the reference electrode, tested in Aptamil solution. **(D)** Calibration curves calculated from experimental results presented in A-C. **(E)** Calibration experiment of K⁺/Na⁺ sensors using the same configuration as in C after 24 h of submersion in Aptamil solution

Ideal Nernstian sensitivity is approximately 59 mV/decade for monovalent ions under perfect equilibrium conditions; however, actual sensor sensitivities often differ due to membrane properties, ion exchange kinetics, and sample matrix effects. The matrix effects of Aptamil help explain the increased sensitivity of the K^+^ sensor in Aptamil, being approximately at Nernst sensitivity at 55.3mV dec^-1^, 1.9x the sensitivity in Tris (27.9 mV dec⁻¹). For Na^+^ in Aptamil, the fully embroidered Na⁺ sensor reached 106.1 mV dec⁻¹, a 1.5 x increase in sensitivity from Tris (70.7 mV dec⁻¹). We suspect that Na⁺ membranes based on Sodium Ionophore X permit faster ion transport and, in complex media like Aptamil, accumulate extra potentials from coupled ion movement and diffusion/junction potentials, producing an apparent super-Nernstian slope ^[59]^. By contrast, valinomycin-based K⁺ membranes are strongly selective for K⁺ over other ions, which makes them less susceptible to these coupled-flux and junction effects, so their responses usually stay close to the Nernst slope ^[51]^. Only hybrid electrode (consisting of PECOTEX as WE) measurements had an OCP that varied linearly with ion activity, further demonstrating the inadequacy of using AgPT exclusively for the electrochemical sensors. We completed all measurements within 1 h of submersion without pretreatment, still maintaining low variability across technical repeats (same sensor for multiple measurements).

Finally, after soaking for 24 h (K⁺: 13 mM KCl; Na⁺: 5 mM NaCl), we assessed post-immersion stability in Aptamil using only sub-immersion concentrations (K⁺: 1–13 mM; Na⁺: 0.3–5 mM) and observed reduced sensitivities for both sensors (**Figure 5E**). The long soak likely caused the ion-selective membranes to become oversaturated. Subsequently, when we tested the sensors over a lower concentration than before (**Figure 5A-C**), the potential outputs decreased in magnitude (K^+^ sensitivity dropped by about 1.4 times; Na^+^ by about 4.3 times), resulting in sub-Nernstian responses (K^+^ sensitivity 40.7 mV dec^-1^; Na^+^ sensitivity 26.67 mVdec^-1^). In this case, we suspect the pre-loading of the membranes limited the ion exchange driving force, resulting in smaller voltage changes than the ideal Nernstian slope ^[60]^. Additionally, the transition from super-Nernstian to sub-Nernstian Na⁺ sensitivity may reflect the loss of additional response enhancements that were present at higher spiking concentrations in the complex matrix ^[60]^. Even with these changes, the sensors still responded with a high degree of linearity to changes in potassium and sodium levels, but we observed increased variability between tests conducted between sensors. To compensate for variability in sensor performance, it is therefore important to calibrate the sensors before each use to maintain accuracy. Despite using a more complex matrix for characterization, the sensor drift over 24 h remained consistent with literature values at approximately 0.28 mV/h ^[61]^.

## 4. Conclusion

The dual thread strategy opens an entirely new range of possibilities for sensing of biophysical and biochemical targets using industrially scalable wearable textile-based sensors. We addressed key bottlenecks in the reliability and durability of textile-based sensors through the integrated platform of the SmartBra. Optimized stitch patterns improved signal quality and wearability: wide satin stitches created continuous surfaces for ion tracking, while zigzag stitches preserved elasticity during deformation. We used redundant re-embroidered connections, which lowered resistance by >38%, and ensured encapsulation protected the circuits against warping during body movement. Our choice of snap-fastener interfaces enabled solder- and adhesive-free connection to flexible electronics, providing robust electrical continuity while allowing easy removal of electronics for washing, repair, or upgrade. Together, these advances directly confront the long-standing challenge of interfacing rigid electronics with conformable garments.

We prioritized off-the-shelf and inexpensive materials (cotton-based threads, AgPT, sports bras, and fasteners) to facilitate real-world adoption by lowering both prototyping and manufacturing costs. We ensured that fabrication relies entirely on CEmb machinery already ubiquitous in textile production, eliminating the need for bespoke tools or processes. With industrial throughput (∼1200 stitches/min), the platform is directly scalable, while the use of familiar apparel ensures comfort and ease of adoption. Together, these strategies position the SmartBra as a realistic candidate for mass manufacturing and everyday deployment, offering a scalable route to personalized, comfortable health monitoring. In the context of maternal and infant health, this integrated sensing garment lays the groundwork for real-time tracking of cardiac activity and breast-milk composition, metrics essential for adaptive nourishment, lactation assessment, and improved infant development outcomes.

The dual thread approach presented in this work also has limitations. First, although PECOTEX offers a cost-effective alternative to AgPT, its conductivity degrades after embroidery and washing due to mechanical stress and material wear. This deterioration can be partially mitigated by tuning embroidery parameters such as stitch density, pattern, and tension, but future work should explore more durable protective coatings, such as thin polyurethane or silicone coatings ^[62]^. Second, the embroidered electrochemical sensors exhibited reduced sensitivity during prolonged exposure to liquids. Integrating protective encapsulation layers such as Nafion could slow down fouling and reduce water ingress, and extend device longevity ^[63]^. Third, our demonstration was conducted in controlled laboratory settings using breast-milk analogues. Real-world performance across diverse users and physiological conditions remains untested, and future work should include clinical validation, multiplexed sensor arrays, and calibration protocols to ensure reliability and user-independent performance.

Looking ahead, the methods presented, especially the SmartBra, pave the way for a new generation of garment-integrated biosensors designed for maternal and neonatal health. For mothers, embroidered electrochemical platforms could be expanded to monitor breast-milk hormones (e.g., cortisol, estradiol) and immune markers (IgA, IgG) alongside electrolytes, offering richer postpartum profiling and earlier detection of lactation or nutritional imbalances ^[64]^. For neonates, the same embroidered architecture could be seamlessly integrated into baby clothing, where conformable, non-removable sensors would ensure continuous monitoring of vital signs and feeding-related biomarkers in a safe, unobtrusive way. Beyond maternal care, the embroidery-compatible approach could be adapted for sweat-based monitoring in athletes to track hydration and electrolyte status, or for military and industrial workers to assess fatigue and heat stress. By embedding electrochemical and physiological sensors directly into garments produced via established embroidery workflows, this platform illustrates how textiles can evolve into scalable diagnostic hubs with capabilities for continuous monitoring within personalized healthcare ecosystems. By continuing to advance the interface between textiles and electronics, we can unlock new avenues in personalized healthcare, bridging the gap between digital health innovation and practical, everyday use.

## 5. Author Contributions

TW performed thread and sensor characterization experiments, designed the sensor and electronics, and performed post-processing work. TW also carried out the ECG experiments and analyzed the resulting data. XL performed the electrochemical experiments and ionophore fabrication and assisted with ECG data collection and photographs. TW wrote the manuscript and produced figures with ASP and HK. MA, JMRF, and LGM assisted with the design of the electrochemical experiments, while SO, HK, MY, and FG assisted with the overall concept of the paper, supervised the research, and revised the manuscript.

## Acknowledgment

We would like to thank EPSRC (EP/G037515/1, EP/L016702/1, Imperial Impact Acceleration Account), Cytiva, Imperial College London, Department of Bioengineering, Gates Foundation (Grand Challenges Explorations scheme under grant number: OPP1212574 and INV-038695) and the US Army (U.S. Army Foreign Technology (and Science) Assessment Support program under grant number: W911QY-20-R-0022), Wellcome Trust (Grant No. 207687/Z/17/Z) and Innovate UK (10004425). TW would like to acknowledge the Skye Foundation for the 2022 Scholarship Award. HK acknowledges the EU Horizon Europe Marie Skłodowska-Curie fellowship (Ref: 101111321) and UKRI MSCA fellowship (EP/Y030273/1) for their support. LGM thanks the European Union’s Horizon 2020 research and innovation program under the Marie Sklodowska-Curie grant agreement no. 101025390. We would also like to thank Noor Khazem for her assistance in operating the Industrial Brother embroidery machine, Beyza Nur Günaydın for the acquisition of SEM micrographs, Ethan Zhou and Cansu Ayabakan for their help with sensor unpicking.

