## Supplementary Information for "Bioelectronic Wearable Sensors Produced by Computerized Embroidery Using a Dual Thread Approach"

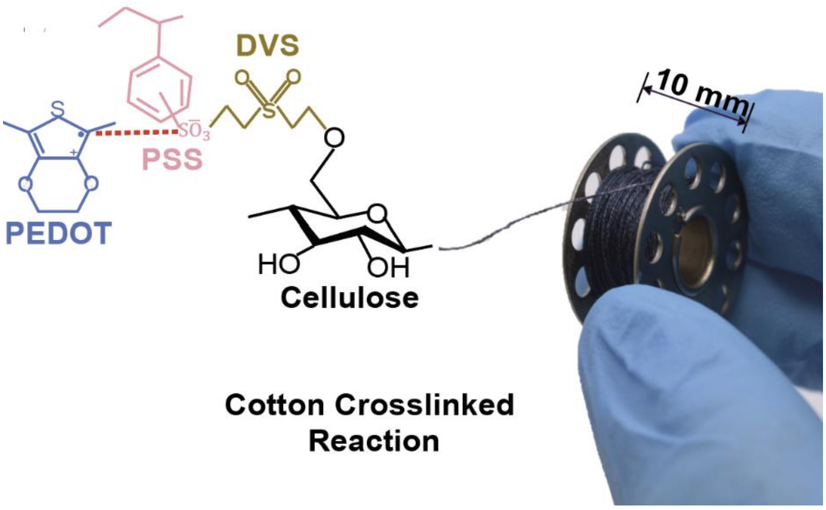


**Figure S1.** Photograph of a manufactured bobbin of PECOTEX demonstrating the crosslinking reaction between the cotton's cellulose and the PEDOT: PSS from the dye.

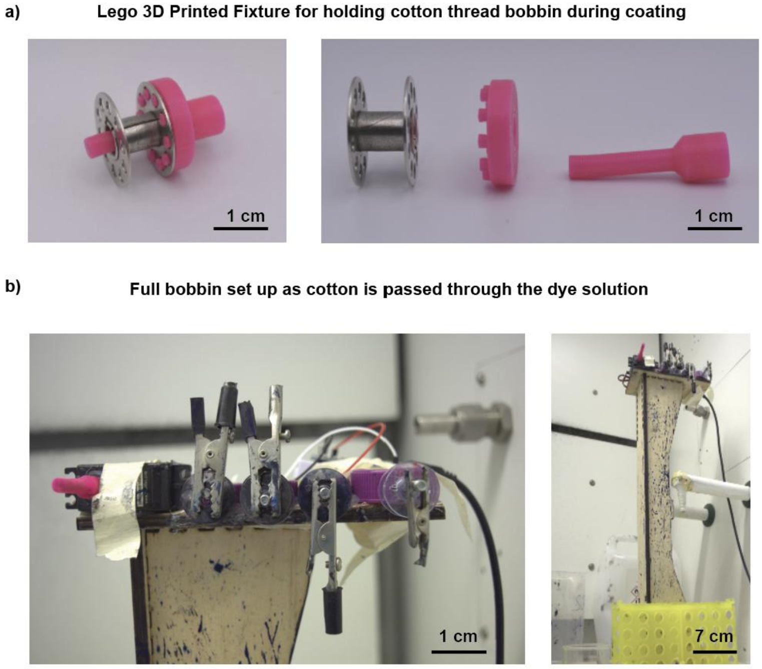


**Figure S2.** PECOTEX producing setup for coating the virgin cotton threads depicting the (a) attachment mechanism for the bobbin and the (b) entire setup.

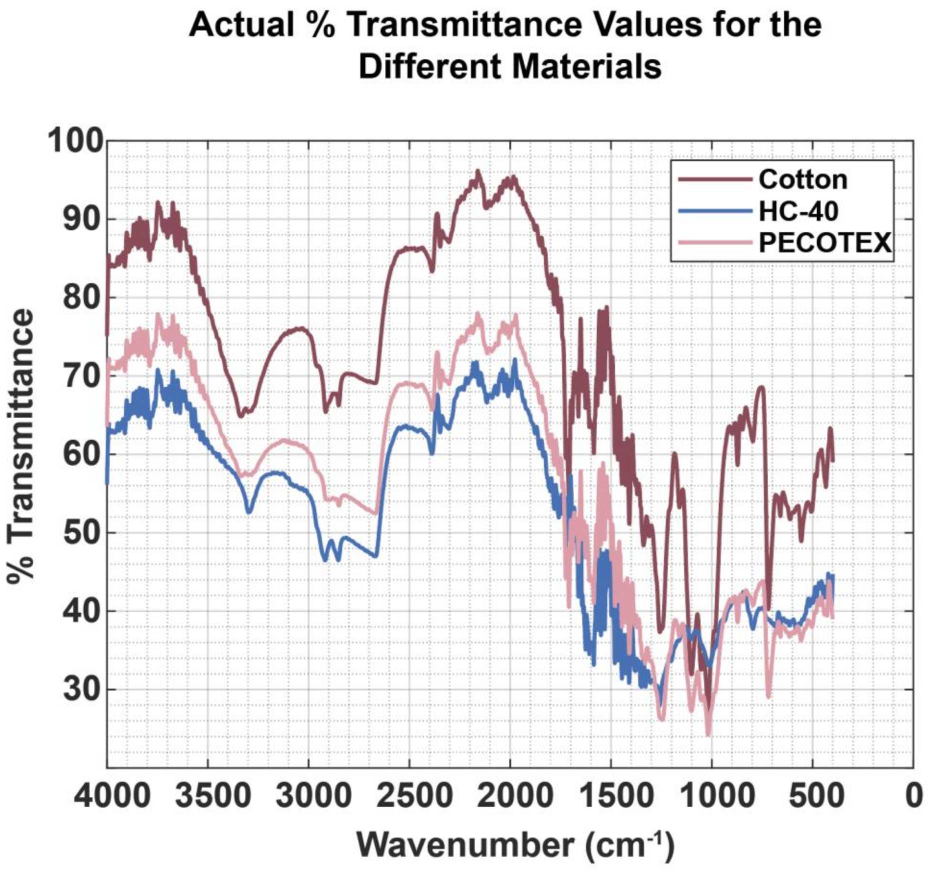


**Figure S3.** The actual % transmittance values for the FTIR-ATR of the different thread types

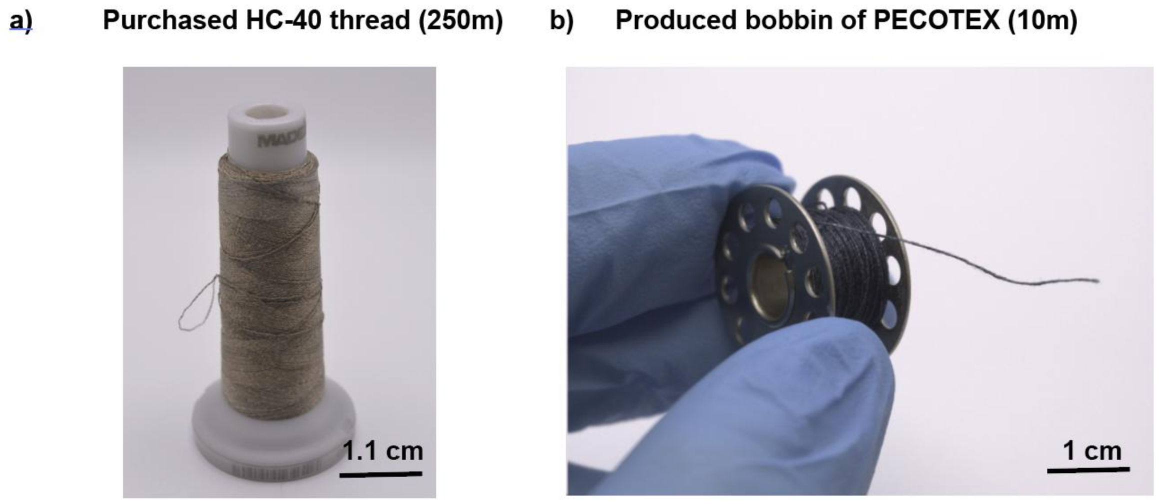


**Figure S4.** (a) Depiction of a purchased spool of HC-40 thread and (b) a produced bobbin of PECOTEX.

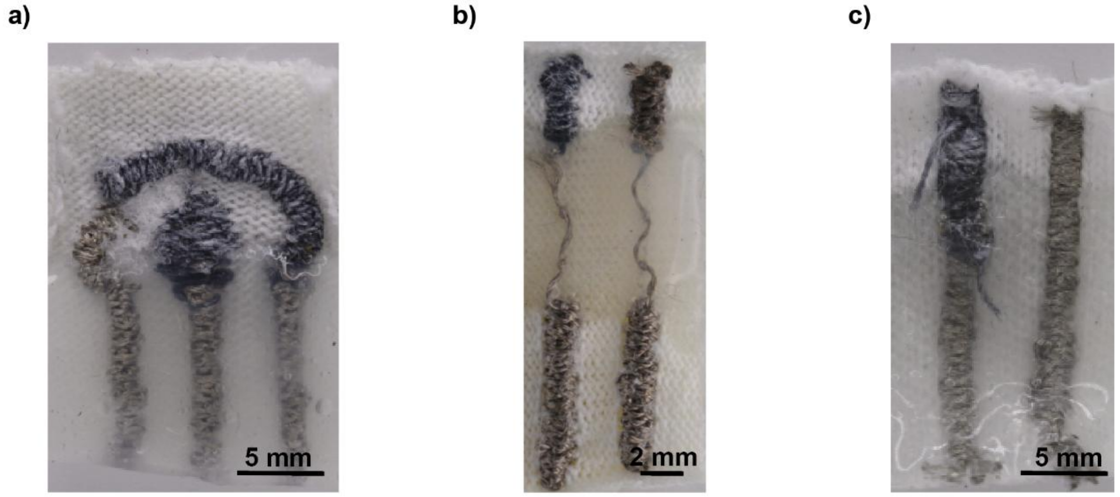


**Figure S5.** The different electrochemical sensors tried with the blue thread being PECOTEX and the silver being HC-40. All sensors have a layer of silicone between the sensor region and the region for crocodile clip (a) Three-pronged electrochemical sensor design in the style of a screen printed electrode with a counter to working electrode width of 1:2; (b) a working and reference electrode with a single thread connecting the sensor region to the crocodile clip region; (c) A smaller design of a two-pronged electrochemical sensor that is fabricated entirely of the satin stitch.

**Table S1.** Embroidery Machine Comparison Between the Household and Industrial Machines

| **Metrics** | **Machine Type** | |
|---|---|---|
| | **Brother PR1050x Industrial Machine** | **HUSQVARNA VIKING's Designer Diamond Royal** |
| Number of Needles | 10 | 1 |
| Computerised Embroidery | Yes | Yes |
| Positioning | Camera based scanning | Manual placement |
| Thread Tension Parameters | Manuel change via thread guide and thread tension knob turning | Set levels via an incremented scale in the setup menu relative to the machine |
| Embroidery Speed | Set levels between 400-1000 stitches / minute | Set levels via an incremented scale on the side of the machine relative to the machine and can be adjusted with a foot pedal |

**Table S2.** Parameter Definition for the Brother Industrial Machine Testing

| **Tension** | **Thread Guide** | **Thread Tension Knob** | **Bobbin Tension** |
|---|---|---|---|
| High | Closed | One Visible Line | Maximum allowed tightness of the machine |
| Medium | Closed | Two Visible Lines | Medium Tightness |
| Low | Open | Red Line Visible | Minimum Tightness the machine will allow |

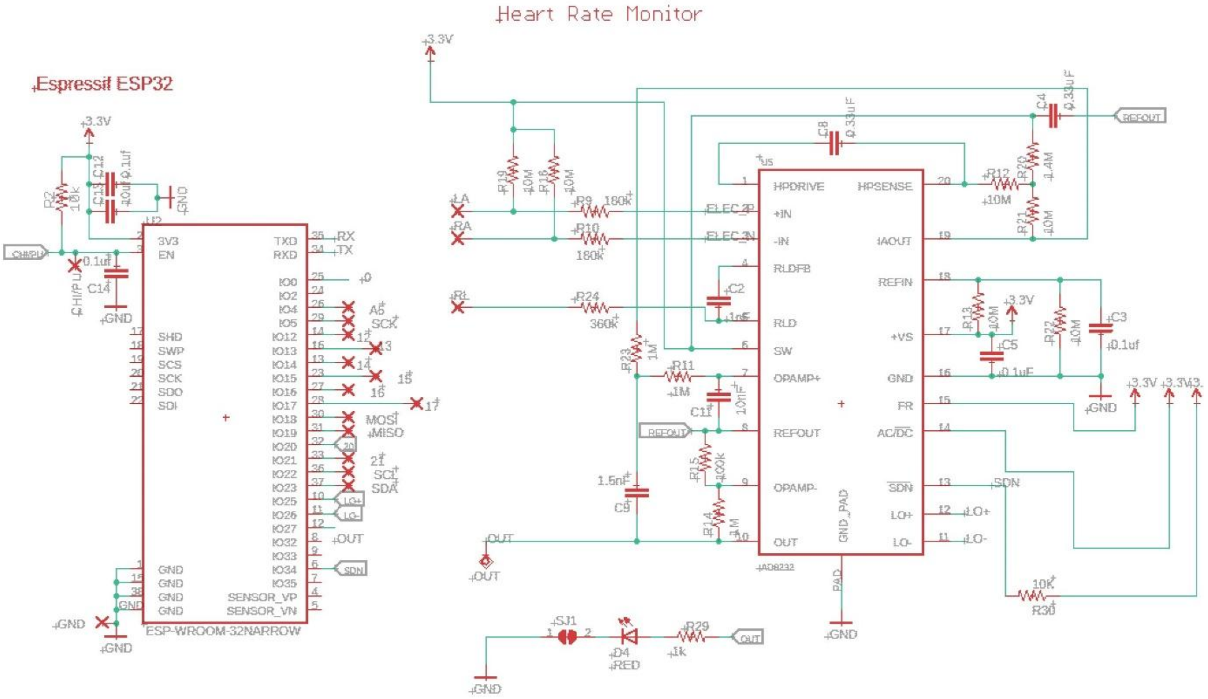


**Figure S6.** Heart rate and microcontroller circuit.

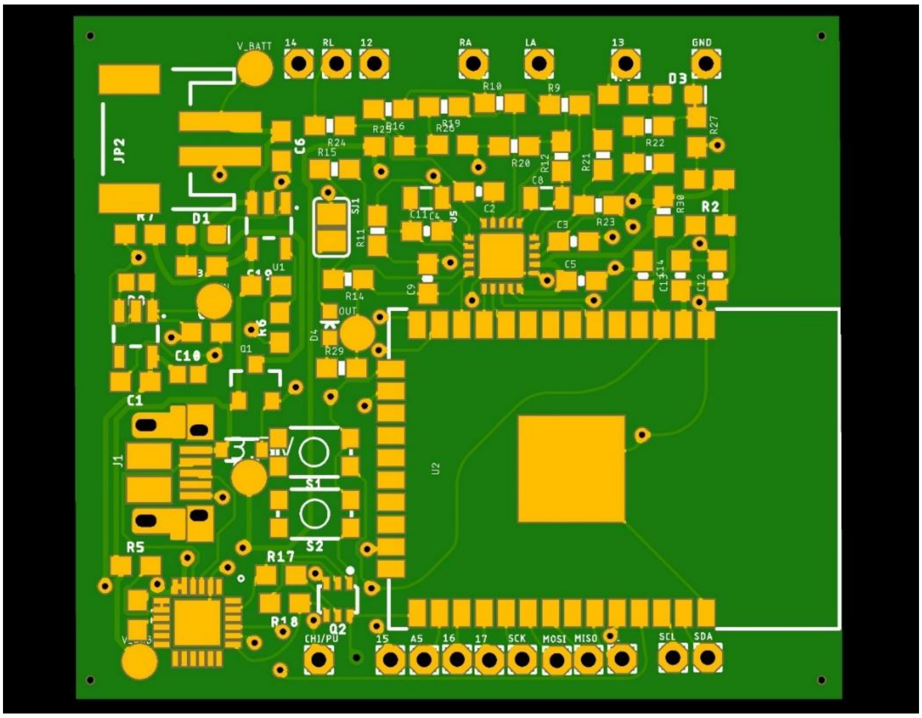


**Figure S7.** Circuit board for component placement, generated in Eagle, with maximal space conservation.

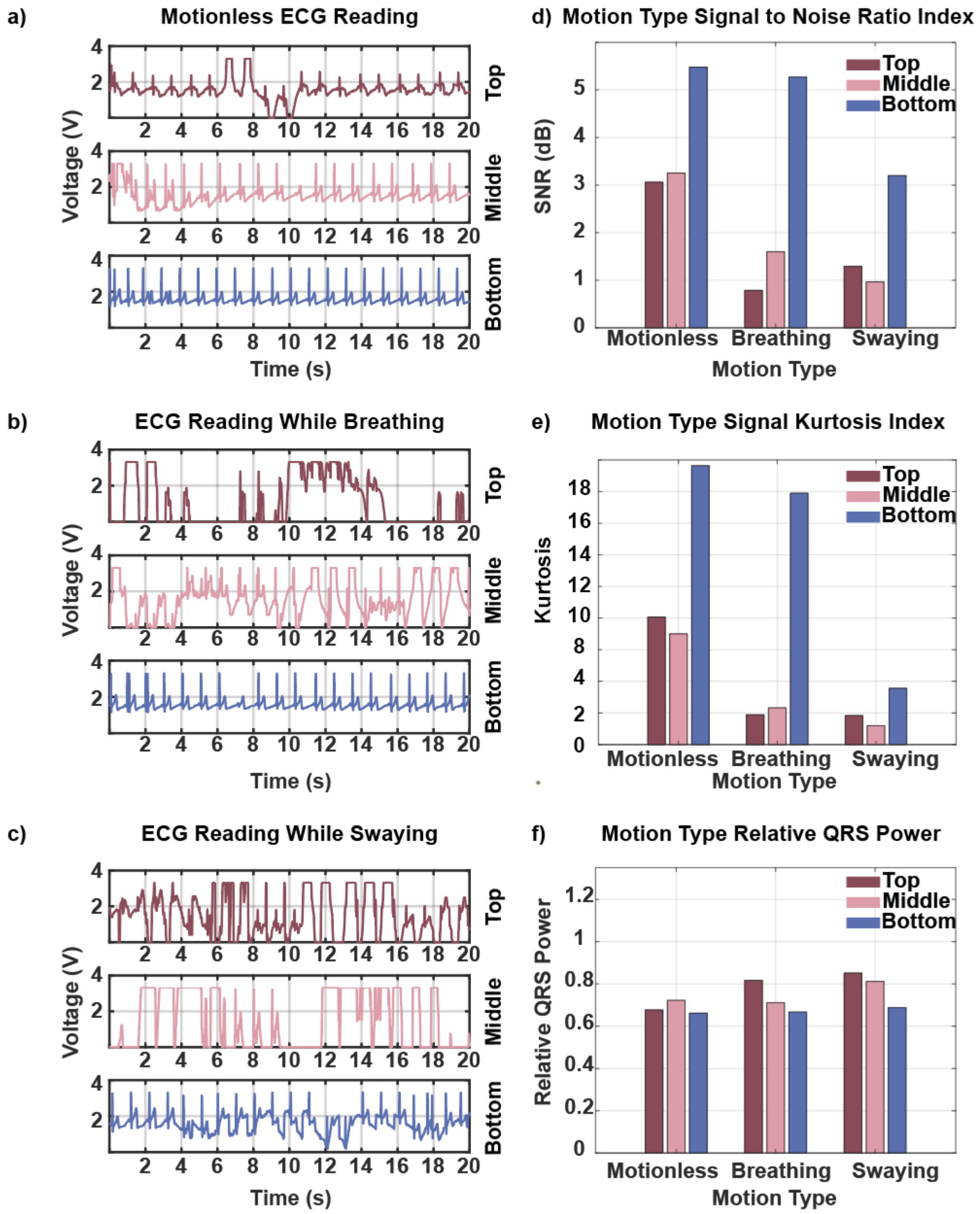


**Figure S8.** (a) The produced ECG from a non-breathing, non-movement position derived from the three locations on the testing bra; (b) The produced ECG while breathing and sitting still; (c) ECG reading while swaying from left to right ; (d) An assessment of the SNR readings for the different electrode positions and motion conditions; (e) The kurtosis values for the different electrode positions and motion conditions; (f) The Relative QRS Power for the different electrode positions and motion condition

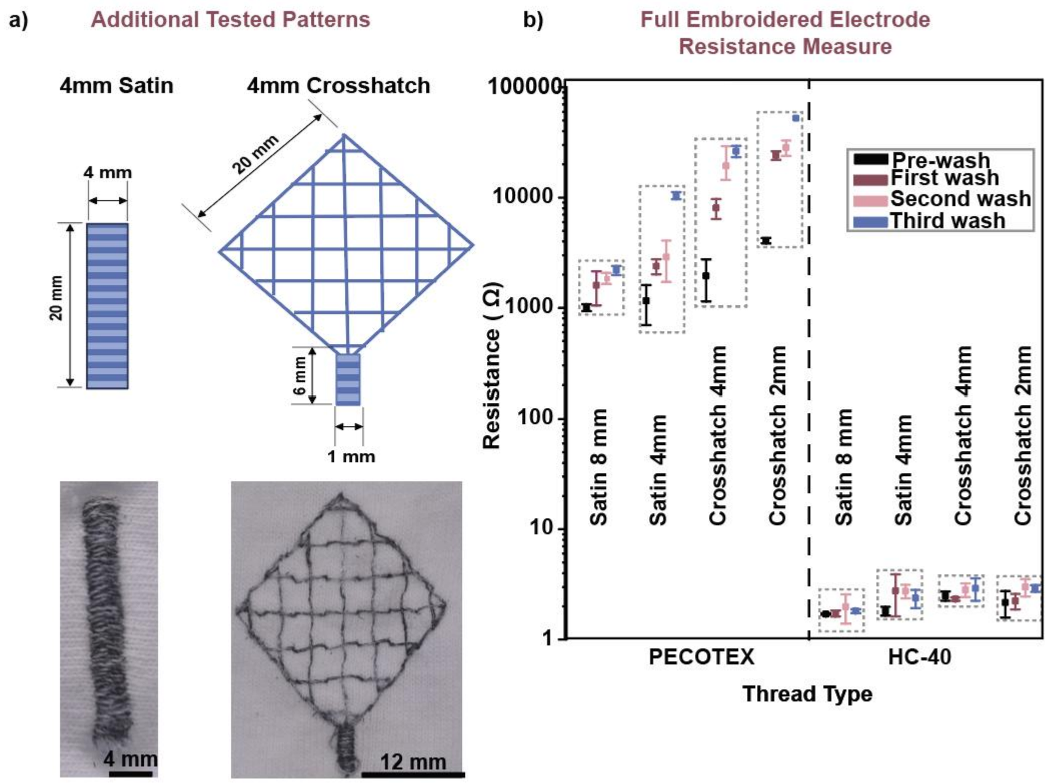


**Figure S9.** (a) Patterns of the satin stitch and crosshatch for the ECG patterns with a different density tested in the washing experiments; (b) Wash tests of the electrode patterns tested

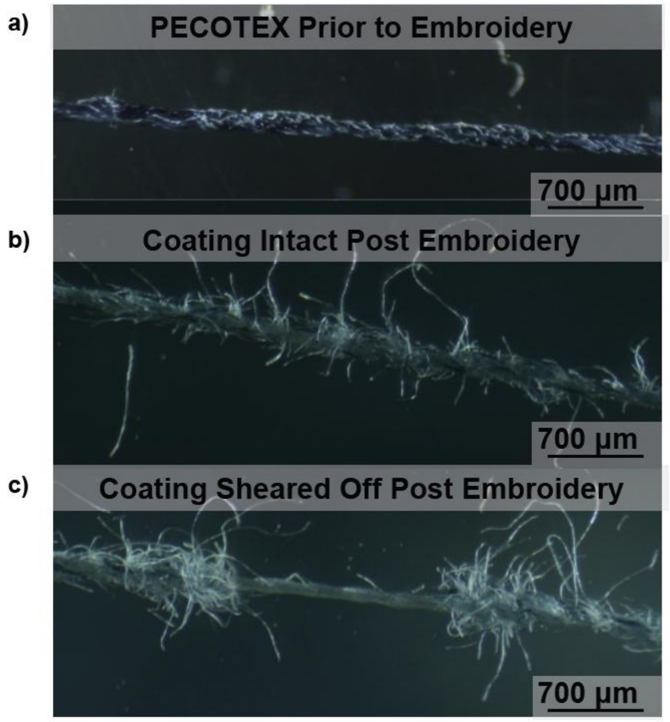


**Figure S10.** Optical imagery of the PECOTEX thread (a) prior to embroidery, and post- embroidery following unstitching. Embroidery damaged the thread with (b) the thread detangling or (c) having the dye completely sheared off.

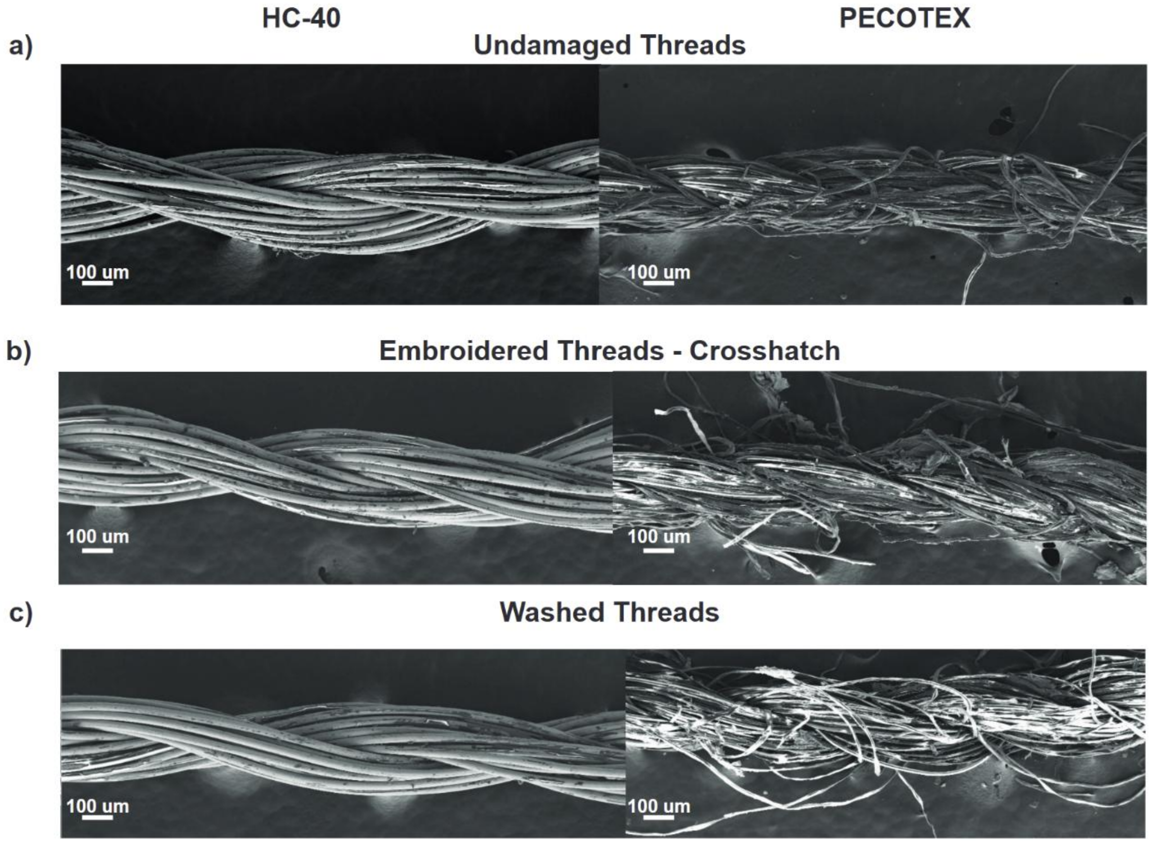


**Figure S11.** SEM Images of both PECOTEC and HC-40 under different conditions: (a) unembroidered, base undamaged threads, (b) threads that have been embroidered with the crosshatch design and subsequently unstitched and (c) threads that have been washed without having been embroidered previously

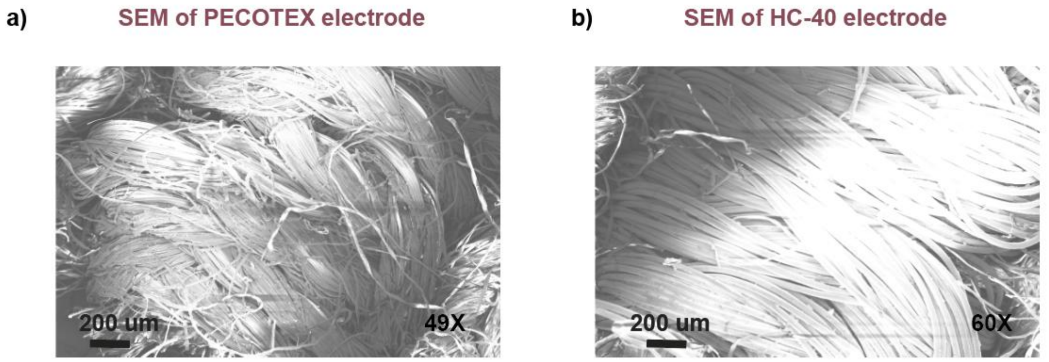


**Figure S12.** SEM Images of the (a) PECOTEC and (b) HC-40 electrodes

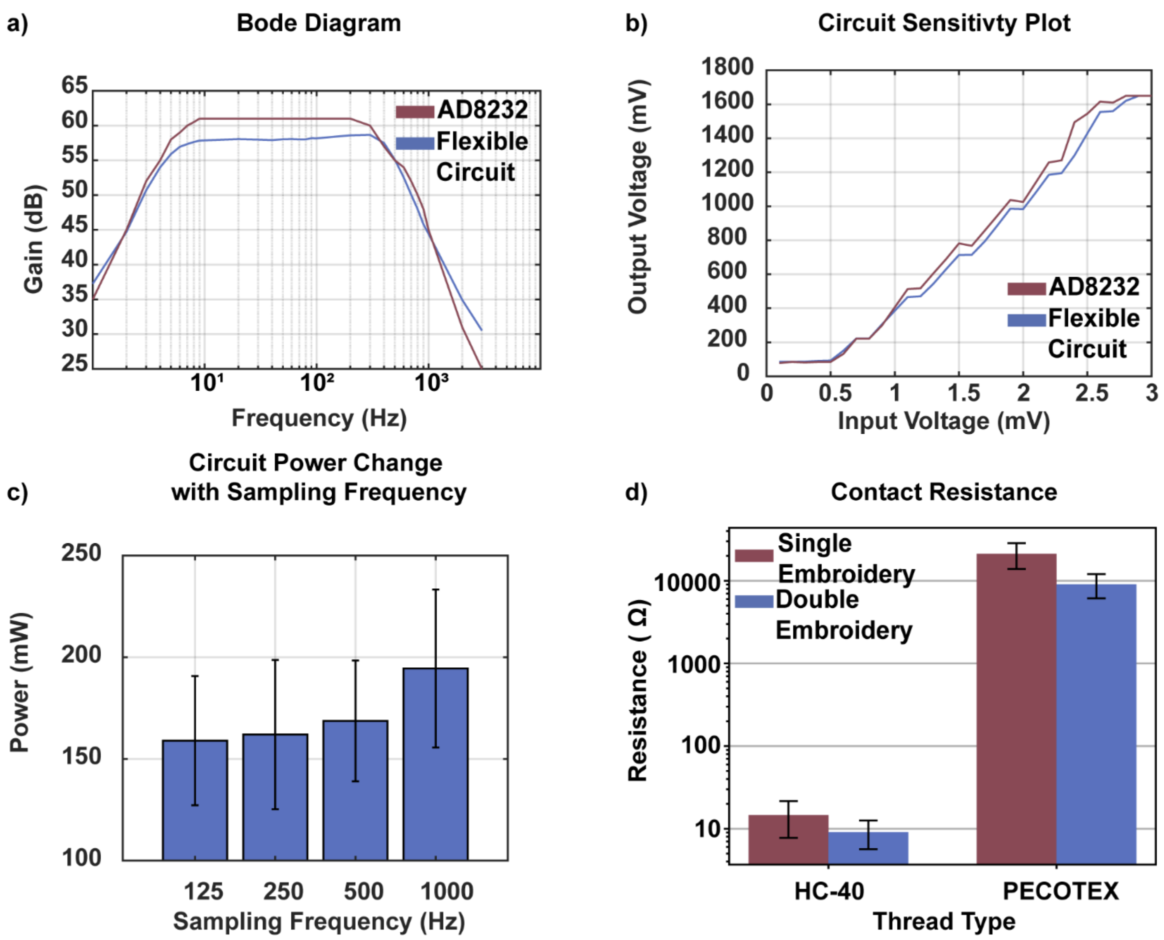


**Figure S13.** (a) The Bode plot comparison between the produced flexible PCB and the AD8232 expected Bode plot in the cardiac monitoring configuration; (b) Comparison of the sensitivity graph of the flexible circuit and the manufactured circuit; (c) Power vs sampling rate determination; (d) Thread and circuit interfacing resistance comparison.

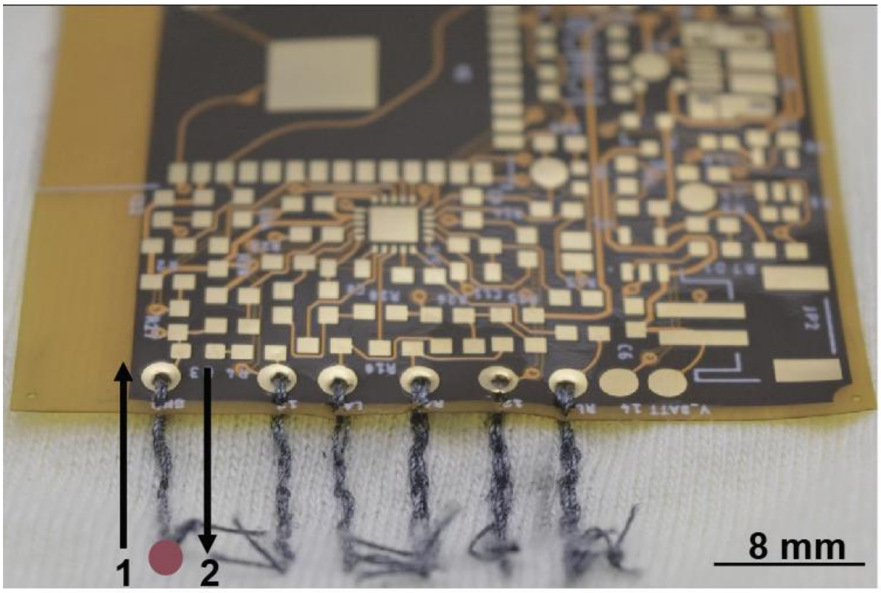


**Figure S14.** The double running stitch interfacing approach begins and terminates at the red circle.

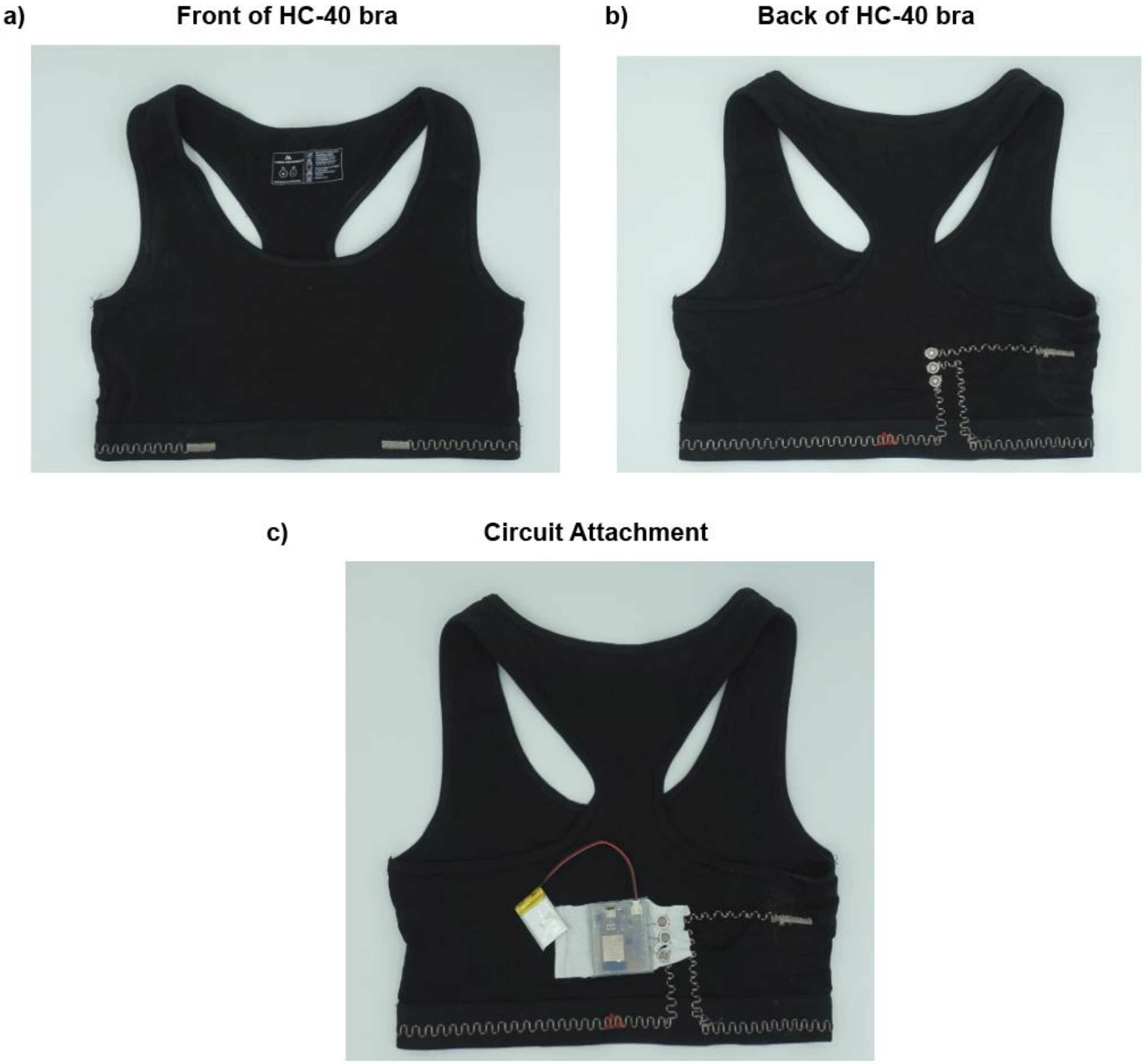


**Figure S15.** HC-40 bra used in the final investigation depicting the (a) front of the bra, (b) back of the bra without the circuit attached and (c) the full circuit attachment with the stud clips.

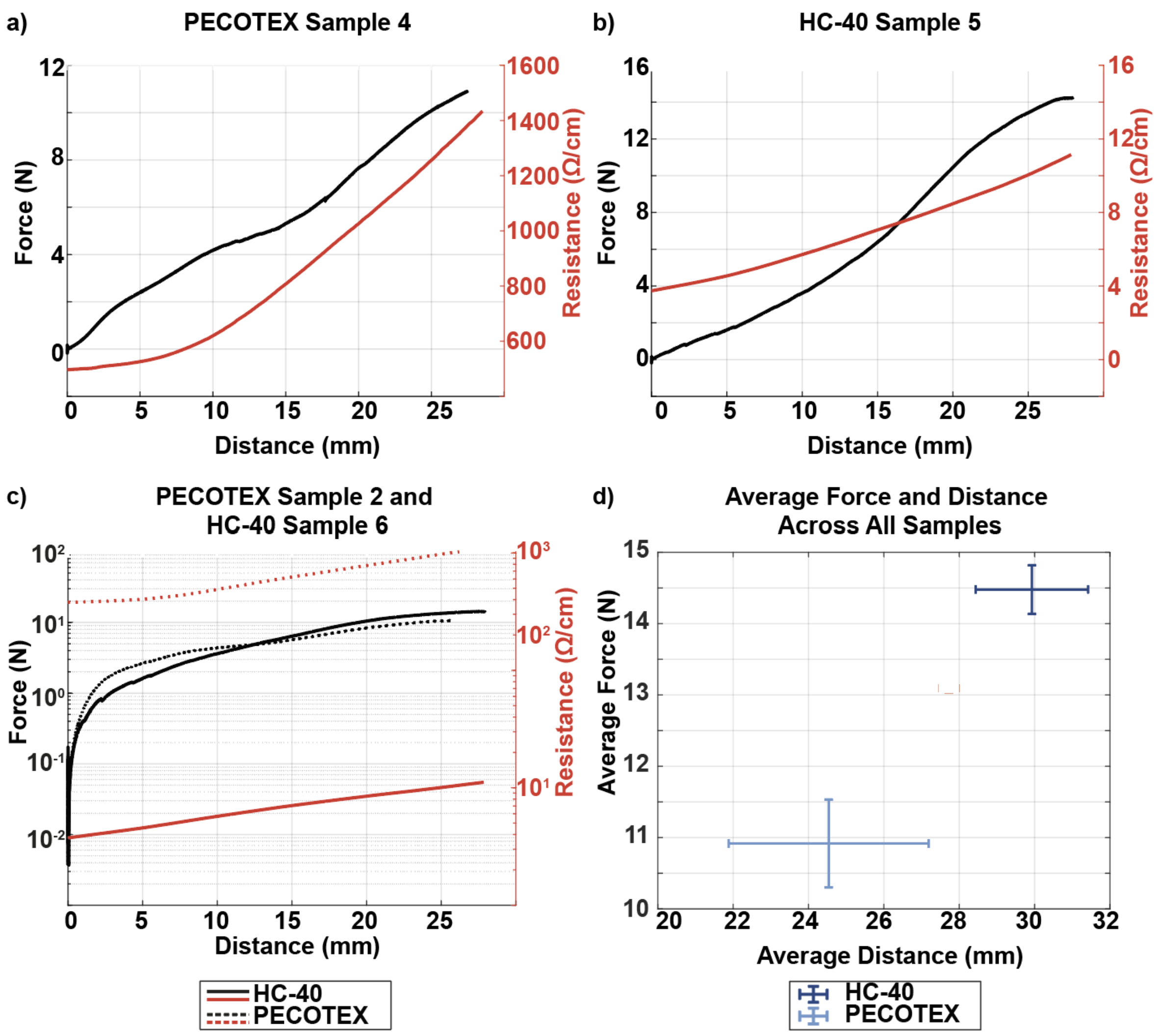


**Figure S16.** (a) PECOTEX and (b) HC-40 thread Force vs Displacement vs Resistance graphs measured on a constant piece of thread with a (c) depiction of their combined change on a single axis; (d) The average force vs distance changes across the 6 samples of each thread type tested.

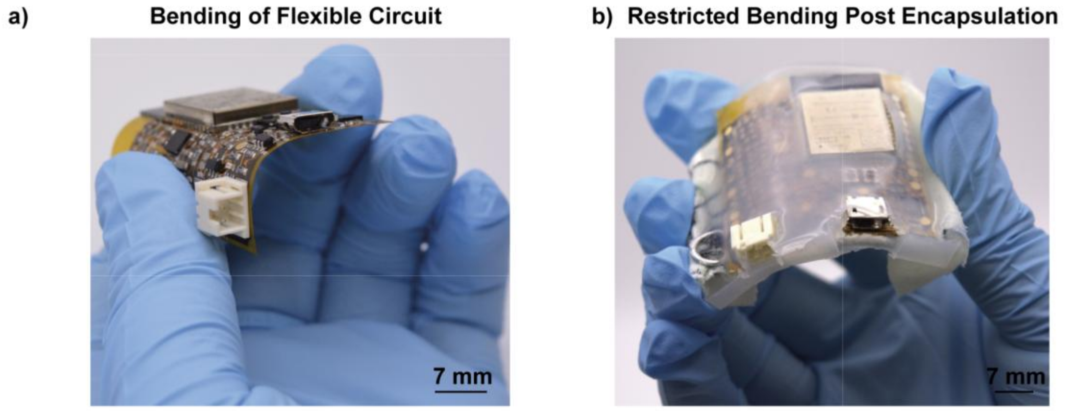


**Figure S17.** (a) The unprotected bending of the flexible circuit pre-encapsulation, as compared to the (b) protected bending of the encapsulated circuit.

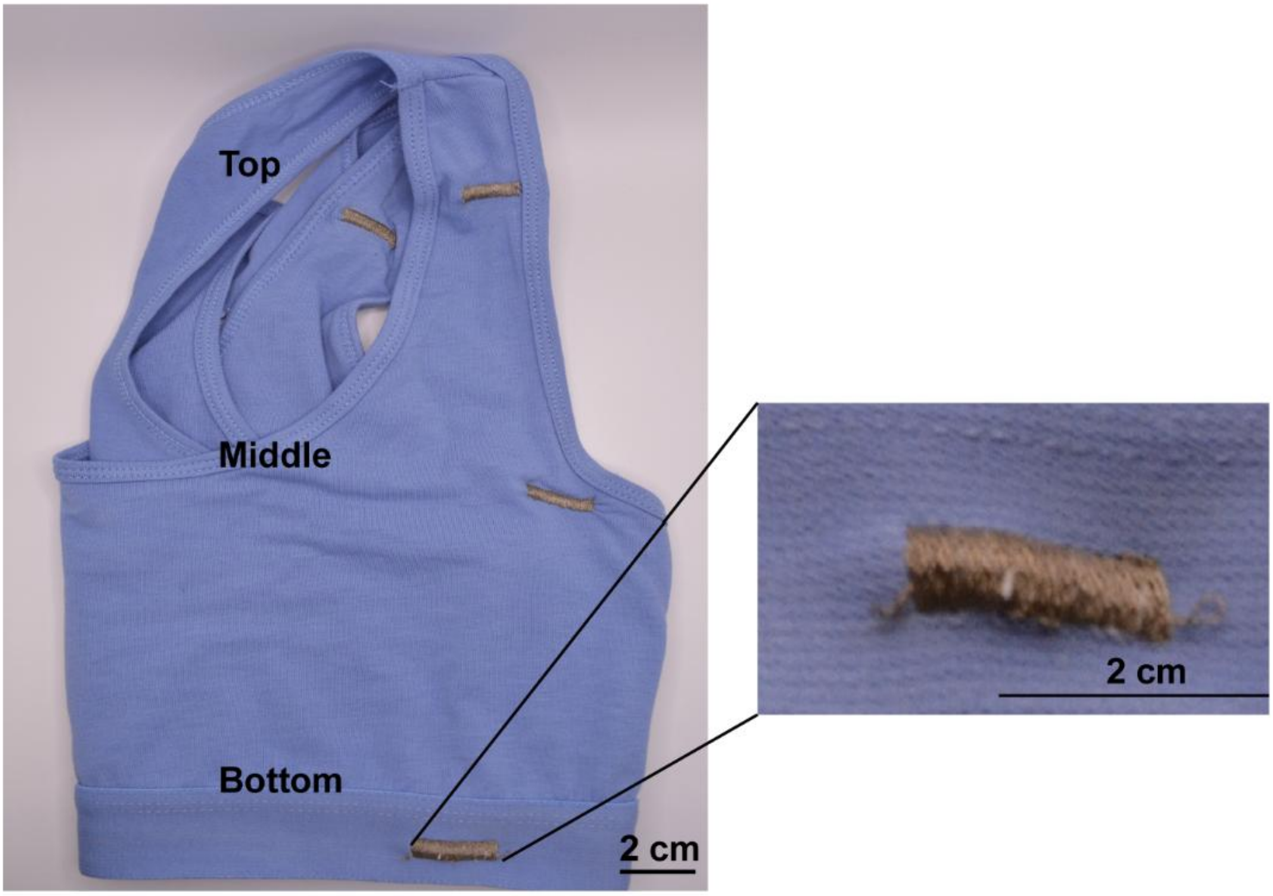


**Figure S18.** Depiction of the produced placement testing bra, which utilised HC-40 thread for prototyping the electrode.
